# Get a new perspective on EEG: Convolutional neural network encoders for parametric t-SNE

**DOI:** 10.1101/2022.12.08.519691

**Authors:** Mats Svantesson, Håkan Olausson, Anders Eklund, Magnus Thordstein

**Affiliations:** Department of Clinical Neurophysiology, University Hospital of Linköping, Sweden; Center for Social and affective Neuroscience, Linköping University, Sweden; Center for Medical Image Science and Visualization, Linköping University, Sweden; Department of Biomedical Engineering, Linköping University, Sweden; Division of Statistics & Machine Learning, Department of Computer and Information Science, Linköping University, Sweden; Department of Biomedical and Clinical Sciences, Linköping University, Sweden

**Keywords:** EEG, Deep learning, Convolutional neural networks, t-SNE, Categories

## Abstract

**Background:** t-distributed stochastic neighbor embedding (t-SNE) is a method for reducing high-dimensional data to a low-dimensional representation and is mostly used for visualizing data. In parametric t-SNE, a neural network learns to reproduce this mapping. When used for EEG analysis, the data is usually first transformed into a set of features, but it is not known which features are optimal.

**New method:** The principle of t-SNE was used to train convolutional neural network (CNN) encoders to learn to produce both a high- and a low-dimensional representation, eliminating the need for feature engineering. A simple neighbor distribution based on ranked distances was used for the high-dimensional representation instead of the traditional normal distribution. To evaluate the method, the Temple University EEG Corpus was used to create three datasets with distinct EEG characters: 1) wakefulness and sleep, 2) interictal epileptiform discharges, and 3) seizure activity.

**Results:** The CNN encoders for the three datasets produced low-dimensional representations of the datasets with a global and local structure that conformed well to the EEG characters and generalized to new data.

**Comparison to existing methods:** Compared to parametric t-SNE for either a short-time Fourier transforms or wavelet representation of the datasets, the developed CNN encoders performed equally well but generally produced a higher degree of clustering.

**Conclusions:** The developed principle is promising and could be further developed to create general tools for exploring relations in EEG data, e.g., visual summaries of recordings, and trends for continuous EEG monitoring. It might also be used to generate features for other types of machine learning.

## 1. Introduction

To understand data, it is necessary to identify important features and from this construct categorizations representing dimensionality reductions. For electroencephalography (EEG) the categorization has historically mainly depended on human experts, but their assessment may suffer from intra- and interrater variability. Furthermore, it is not known if the categories are the most optimal. When developing algorithms or machine learning to analyze EEG, the data often need to be transformed into a set of features, e.g., a limited set of numbers related to the frequency content. However, important information may be lost in this transformation. Hence, developing methods that discover relevant EEG features and how these are related through a self-supervised process may provide a new perspective on traditional EEG categories and possibly new insights. To this end, in the work presented here, a deep learning approach for adapting parametric t-distributed stochastic neighbor embedding (t-SNE) to EEG analysis was taken. This article has been written with both clinical neurophysiologists and data scientists in mind. Many of the technical details have therefore been placed in appendices.

t-SNE is a method primarily developed for visualizing high-dimensional data by mapping it to a low-dimensional space (van der Maaten and Hinton, 2008). The result is usually clustering of similar data in the low-dimensional representation and relations in the data can then be identified by visual inspection and comparisons with the original data (Fig. 1.).

**Fig. 1.**
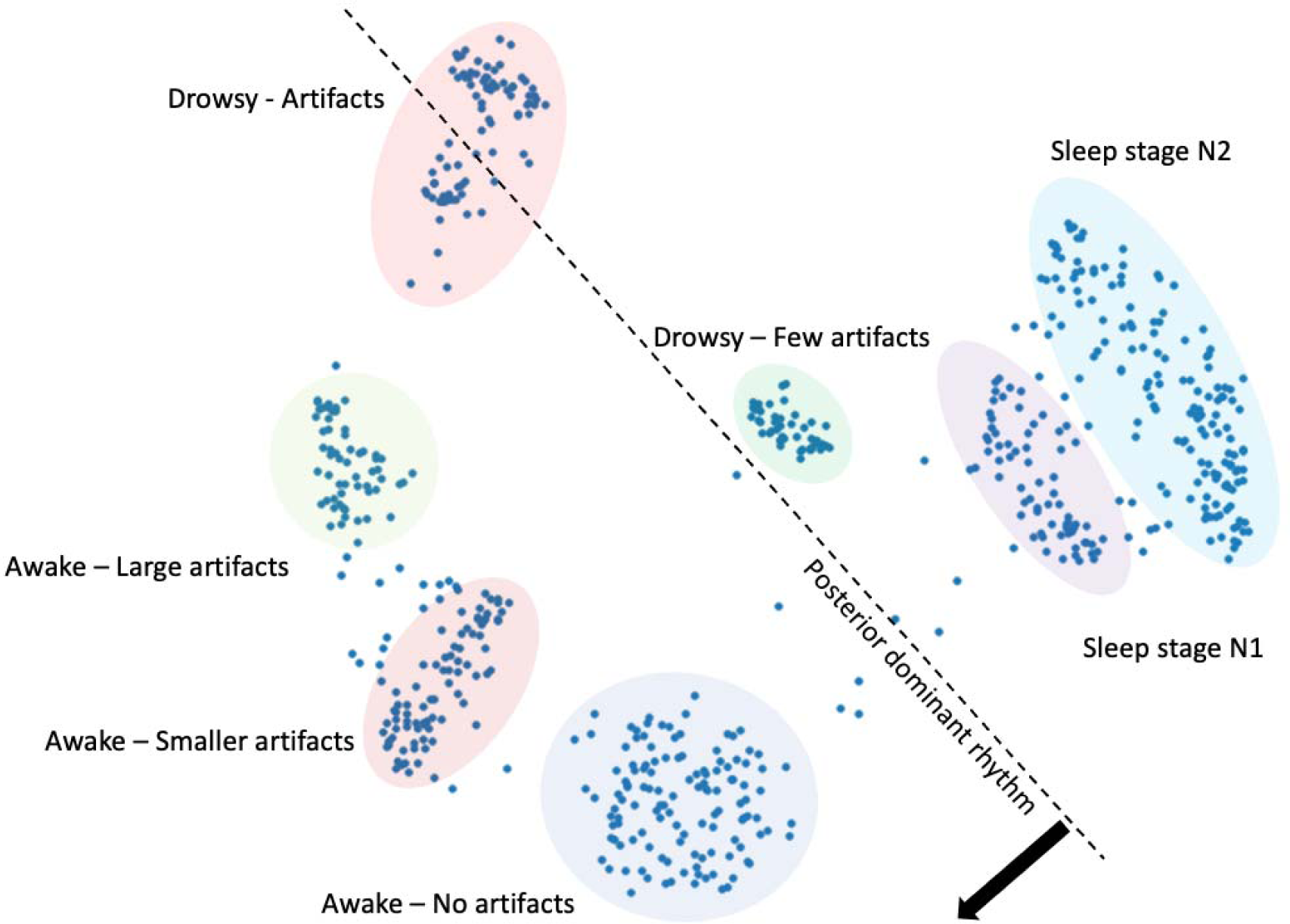
Low-dimensional representation of a 21-channel EEG of 22 minutes duration, where each dot represents 2 s of EEG. The figure was generated by a convolutional neural network trained using the principle described in this article which is based on t-SNE. With a sampling frequency of 250 Hz of the raw EEG, this represents a reduction from 6,930,000 to 1,320 data values. A comparison with the original EEG was made and the structure of the low-dimensional representation is indicated in the figure.

t-SNE is based on pairwise matching of the probabilities that data examples are neighbors in both the high- and the low-dimensional space. The high-dimensional probability distribution is multivariate normal and the low-dimensional is a multivariate t-distribution. Optimization is accomplished by gradient descent. The original implementation does not create a model for the mapping from high- to low-dimensional representation. (The method is described in some more detail in Appendix B.1). The original t-SNE is used for visualization of results in several EEG studies (Jing et al., 2018; Kinney-Lang et al., 2018; Al-Fahad et al., 2020; Ravi et al., 2020; Suetani and Kitajo, 2020; Idowo et al., 2021; Jeon et al., 2021; Kottlarz et al., 2121; Malafeev et al., 2121; Georg et al., 2022). In a couple of other studies, the method is used for feature extraction from EEG (Ma et al., 2021; Yu et al., 2022).

Parametric t-SNE is a variation where the dimensional reduction is learned by a neural network (van der Maaten, 2009), thus creating a model for the mapping (Fig. 3A.). In the original implementation restricted Boltzmann machines are pretrained using greedy layer-wise training and then finetuned using the principle of t-SNE. Li et al. (2016) and Xu et al. (2020) use parametric t-SNE as a step to extract features of motor imagery EEG data and evaluation by support vector machine classifiers compares favorably to other feature extraction methods.

Substantial preprocessing is usually required to extract features from EEG. Jing et al. (2018) use frequency features in combination with several other statistical or nonlinear measures. Suetani and Kitajo (2020) employed frequency features in a modified version of t-SNE using a beta-divergence as distance measure. Kottlarz et al. (2021) use t-SNE to compare frequency features and features created using ordinal pattern analysis. Li et al. (2016) use wavelet generated features for parametric t-SNE, and Xu et al. (2020) extend this by combining wavelet features with the original EEG and frequency band features, all processed further through common spatial pattern filtering. Other researchers create features using tensor decomposition (Kinney-Lang et al., 2018), connectivity measures (Al-Fahad et al., 2019), and several extracts features from latent spaces of neural networks (Ravi et al., 2020; Idowo et al., 2021; Jeon et al., 2021; Malafeev et al., 2021). Exceptions to the above are Ma et al. (2021), Georg et al. (2022), and Yu et al. (2022) who apply t-SNE to raw EEG to create features. It is difficult to assess how the different approaches compare in terms of performance and none of them can be regarded as a standard approach.

An important property when using a clustering method for visual analysis is to, not only produce an order according to similarities of data examples, but also to solve the *crowding problem* (Fig. 2A) and produce distinct separated clusters (Fig. 2C). When working with unannotated data, this is essential to be able to identify potential categories (Fig. 2).

**Fig. 2.**
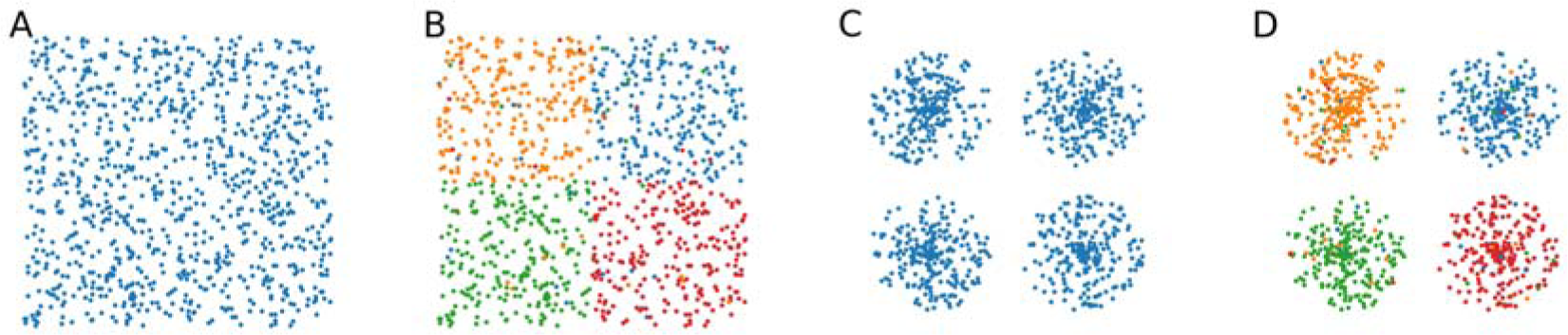
Illustration of clustering using unannotated or annotated data. In A there is no separation of the categories, which can only be seen when using color coding according to an annotation as in B. In C it is easy to see four potential categories, which is confirmed in D when using color coding.

The difference between the statistical distributions for the high- and low-dimensional representation was introduced in t-SNE to solve the crowding problem (van der Maaten and Hinton, 2008), but the data and their high-dimensional representation will of course also affect how well categories are separated by the method. A high-dimensional representation similar to how an expert view EEG might be preferable. Here, the identification of a high-level feature such as interictal epileptiform discharges (IEDs) is regarded as important whereas their time location may be less so. Using the Euclidean distance of the raw data to compare visually similar examples could result in them being assessed as different if important waveforms are located at different positions. Furthermore, if the waveforms only constitute a small part of the signals, they may not have large enough impact on the measure. A property often claimed regarding convolutional neural networks (CNNs) is the location invariant detection of features (LeCun et al., 2015). Therefore, using a CNN with appropriate fields of view to convert the EEG into a set of high-level features seem like a possible solution but the question of how to train the network need to be addressed. Training the networks in classification tasks may be an alternative (Ravi et al., 2020; Idowo et al., 2021; Malafeev et al., 2021), but this may put too much emphasis on features specific to the classification task and the dataset and involves supervised learning.

The objective of this work was to further improve parametric t-SNE for EEG visualization. The main concept was to eliminate the need for extensive preprocessing and feature engineering by instead training CNNs using the principle of t-SNE to match a neighbor structure in its latent space to one in its output space (Fig. 3B.).

**Fig. 3.**
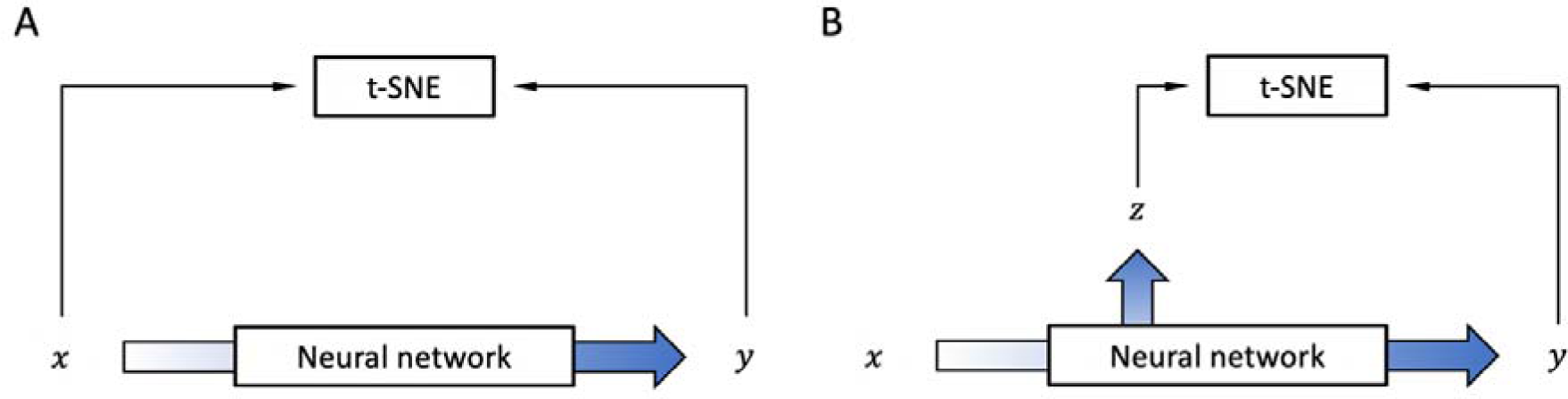
Comparison of the original parametric t-SNE and the suggested method of this work. Both methods produce a low-dimensional representation of the high-dimensional data using a neural network. (A) The original parametric t-SNE. The input is usually a preprocessed version of the EEG. The optimal way to preprocess the EEG is not known. (B) The suggested method. The input x is the raw EEG data and a new high-dimensional representation of the data is extracted from the latent space of the network. An advantage with this approach is that the preprocessing step necessary in (A) is eliminated.

## 2. Materials and methods

### 2.1. EEG data

#### 2.1.1. TUH EEG Corpus

All EEG data used in the study was retrieved from the large, public database of the Temple University Hospital, Philadelphia—the TUH EEG Corpus (Obeid and Picone, 2016). The database contains a mixture of normal and pathological EEGs, and the raw data are unfiltered. The recording electrodes are positioned according to the international 10-20 system (Jasper, 1958). The TUH EEG Corpus (v1.1.0) with average reference was used (previously downloaded in the context of other projects during 17–21 January 2019).

#### 2.1.4. Ethical considerations

The TUH EEG Corpus was created following the HIPAA Privacy Rule (Obeid and Picone, 2016). No personal subject information is available and there is no way of tracing the data back to the subjects. It was therefore not deemed necessary to seek further approval for this study.

#### 2.1.5. Extracted data

From the TUH EEG Corpus three datasets of examples of 1 s duration were created: 1) wakefulness and sleep (24,647 examples of wakefulness and 14,450 examples of sleep), 2) IEDs (4,384 examples containing IEDs and 7,944 examples without IEDs), and 3) seizure activity (2,549 examples of seizure activity and 7,467 examples without seizure activity). The data extraction and annotation processes are described in more detail in Appendix A. Each dataset was split into approximately 75% for training and 25 % for testing. For dataset 1, the test data consisted of new subjects, for dataset 2 and 3, the test data consisted of new examples from the same subjects that were used for training. The standard electrodes of the international 10– 20-system were used: Fp1, F7, T3, T5, Fp2, F8, T4, T6, F3, C3, P3, O1, F4, C4, P4, O2, Fz, Cz, Pz, A1, and A2 (this is also the channel order used in the analyses). Average reference was used.

#### 2.1.6. Preprocessing

The data were band pass filtered between 1 and 40 Hz using Butterworth zero phase filtering. The data were normalized to keep values mostly varying within the interval -1 to 1 by dividing by the 99^th^ percentile of the absolute amplitude of the training data.

### 2.2. Encoders

Details regarding the network architecture, hyperparameters, and training of the encoders are given in Appendix C.

#### 2.2.1. Modification of t-SNE

As described in the introduction and Appendix B.1, t-SNE is based on matching probabilities of data examples being neighbors in a high- and a low-dimensional representation of the data. The suggested approach uses a high-dimensional representation of the EEG data that was generated by the CNN encoders. This means that the representations changed during training, and it was necessary to generate them for every batch of training data to compute the corresponding probability distribution. To simplify the computations, a simple distribution based on ranked distances was used instead of a normal distribution. To compute the former distribution, the distances between examples were ranked in increasing order and the u closest relative to each example were regarded as-, and therefore given a high probability of being neighbors, while all others were given a low probability. The parameter *n* thus had a similar function as perplexity has in the original t-SNE. See Appendix B.2 for a more detailed description.

#### 2.2.2. CNN encoder architecture

The CNN encoders consisted of two parts, one part that learned to produce a high-dimensional representation *z_i_* and another part that in turn mapped this to a low-dimensional representation *y_i_* (Fig. 4).

**Fig. 4.**
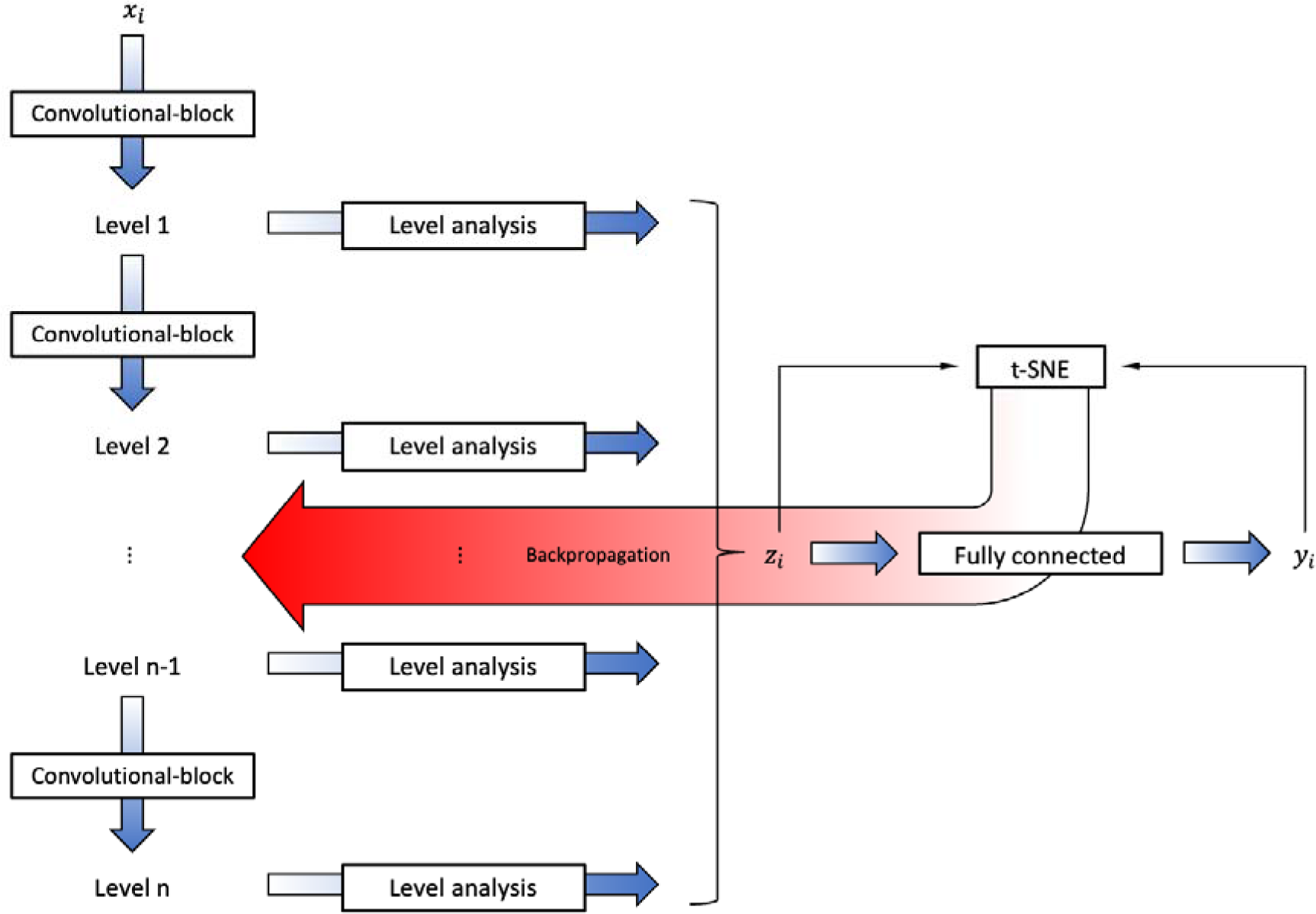
Illustration of the CNN encoder architecture. To the far left a series of blocks of convolutional and max pooling layers generate levels with increasing field of view of the EEG ( ). Each level is then processed by further convolution, max pooling, and then a small fully connected layer to produce a set of values which together form the latent representation of A fully connected network then maps into a low-dimensional representation . The principle of t-SNE is used to compute a loss from which the gradient is determined in the backpropagation.

The first part had a structure that created levels with different field of view of the data, then extracted values that reflected the character of each level. The values all together then formed the high-dimensional representation of the data. This was implemented as a series of convolutional and max pooling blocks, the latter also down sampling the data, thereby producing a set of levels. The levels were then processed by a combination of a convolutional layer, max pooling, small fully connected layers, and the resulting features from all levels were concatenated to form the high-dimensional representation. EEG-channels were analyzed separately in the first part.

The second part was implemented as a series of fully connected layers where the last layer had two nodes that produced the low-dimensional representation.

For further information on the architecture, see Appendix C.

### 2.3. Evaluation

CNN encoders were trained for the three datasets. Training was performed using a batch size of 500 and 5 neighbors per batch (Appendix B.4).

#### 2.3.1. Comparative methods

Comparisons for the developed method were made with in-house implementations of parametric t-SNE using two different feature types: short-time Fourier transforms (STFT) and continuous wavelet transforms (CWT). These implementations are described in more detail in Appendix D. In short, the STFT approach produces a set of values corresponding to frequency bands for short time intervals, in this case 1 s and 30 bands in the range ∼1–30 Hz. This resulted in 630 values per example. CWT also produce values corresponding to frequency bands for each EEG channel. 29 bands with most of the frequency content below 30 Hz were used and the following statistics for each band were calculated: mean, standard deviation, minimum and maximum value. This resulted in 2,436 values per example. Training was performed using a batch size of 500 and a perplexity of 5 for both feature alternatives.

#### 2.3.2. Quantitative measures

The developed method and parametric t-SNE were here used to produce two-dimensional representations of EEG data. Quantifying the quality of an image appears difficult, but two types of measures that reflect how the categories were separated were produced in addition to the resulting images by means of support vector machines (SVMs) and k-means clustering.

The three datasets presented in section 2.1.5 each consisted of two categories. Using 0 and 1 as labels for the categories of each dataset, SVMs were fitted to the resulting low-dimensional representations of the training data and evaluated on both the training and test data to produce quantitative measures of the separability of the categories and generalization of the models. Linear separability would indicate that the global structure is simple. Since t-SNE produces a nonlinear projection of the data with an element of chance, linear separability may not be good even though the categories are well separated in distinct clusters. Separate SVMs were therefore fitted to the data using either a linear or a radial basis function (RBF) kernel. The number of examples was different for each category and a kappa score 1< (Cohen, 1960) was used instead of accuracy to adjust for chance results. The resulting classifications were also compared using a chi-square test. If there was a significant difference, post hoc analysis was performed comparing the best classification with the two others individually and applying Bonferroni adjusted p-values.

K-means clustering was used to produce a measure of the ability to generate distinct clusters of the data by each method. Tests from 2 up to 50 clusters were performed and the number *C* of clusters giving the best silhouette score (Rousseeuw, 1987) was taken as the measure.

For all measures, higher values were regarded as better. P-values below 0.05 were considered as significant.

### 2.4. Software and hardware

The CNN encoders were developed in Python (version 3.7.13) using the API Keras (version 2.8.0) and programming module TensorFlow (version 2.8.0). The Python library ‘pyEDFlib’ (version 0.1.22) (Nahrstaedt and Lee-Messer, 2017) was used to extract EEG data. Filters were implemented using the ‘scipy.signal’ library (Virtanen et al., 2020). Scikit-learn (Pedregosa et al., 2011) was used for support vector machines and k-means clustering.

Four different computers (running Ubuntu) were used, equipped with 64–128 GB of RAM and either one or two Nvidia graphics cards among: Quadro P5000, Quadro P5000 RTX, Quadro P6000, Titan RTX, or Titan X.

## 3. Results

### 3.1. Sleep-wake

All methods produced high kappa scores (Tab. 1), but overall, the CWT encoder produced the highest scores, where the scores were significantly highest when using a linear kernel. When training with an RBF kernel the CNN encoder scored highest, but for the test data, the STFT and the CWT encoders scored equal and significantly higher than the CNN encoder.

**Tab. 1.**
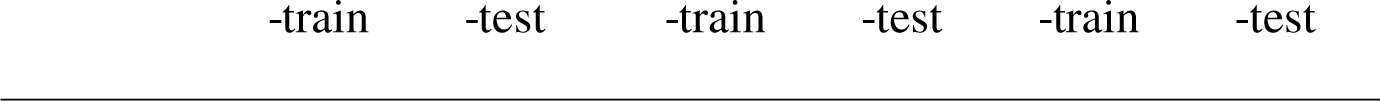
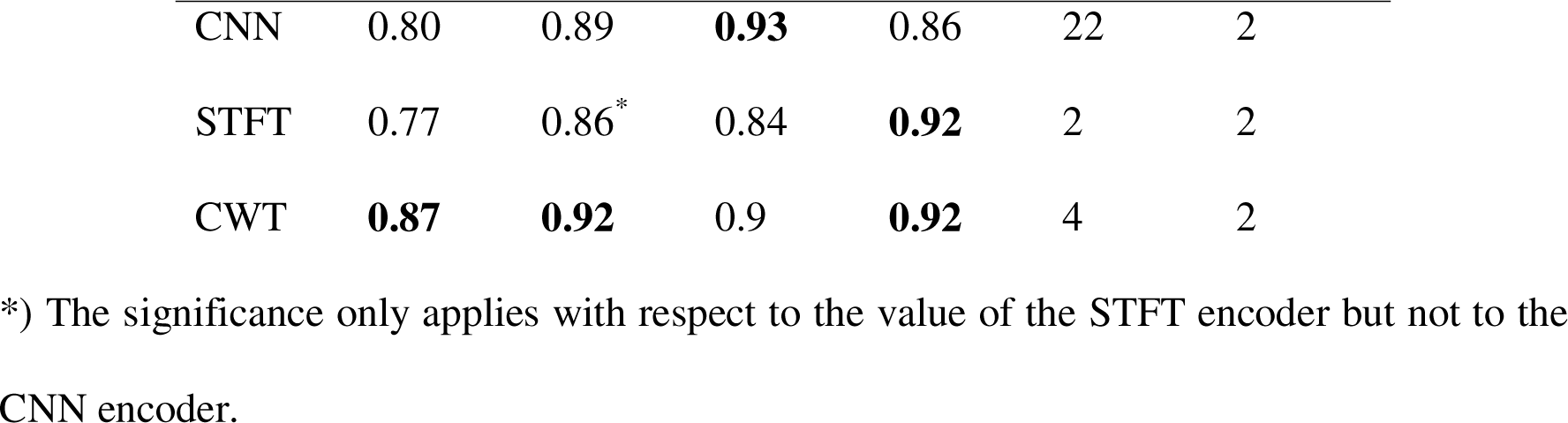
Quantitative measures for the sleep-wake dataset. Significantly highest values in a column are in bold (chi-square test); are kappa values for SVM, indicates a linear kernel and a radialis basis function kernel; indicates the number of clusters by k-means.

|  |  |  |  |  |  |  |
| --- | --- | --- | --- | --- | --- | --- |
| CNN | 0.80 | 0.89 | **0.93** | 0.86 | 22 | 2 |
| STFT | 0.77 | 0.86* | 0.84 | **0.92** | 2 | 2 |
| CWT | **0.87** | **0.92** | 0.9 | **0.92** | 4 | 2 |

\*) The significance only applies with respect to the value of the STFT encoder but not to the CNN encoder.

The CNN encoder produced the highest clustering score for the training data (Tab.1), where there visually was an apparent difference compared to the other methods (Fig. 5B, 5E, and 5H), but for the test data there were few clusters (Fig. 5C), so it generalized but with less substructure. For all methods the global structure was simple, with two main areas containing most of the respective categories. This was in line with the relatively high kappa scores when using a linear kernel.

**Fig. 5.**
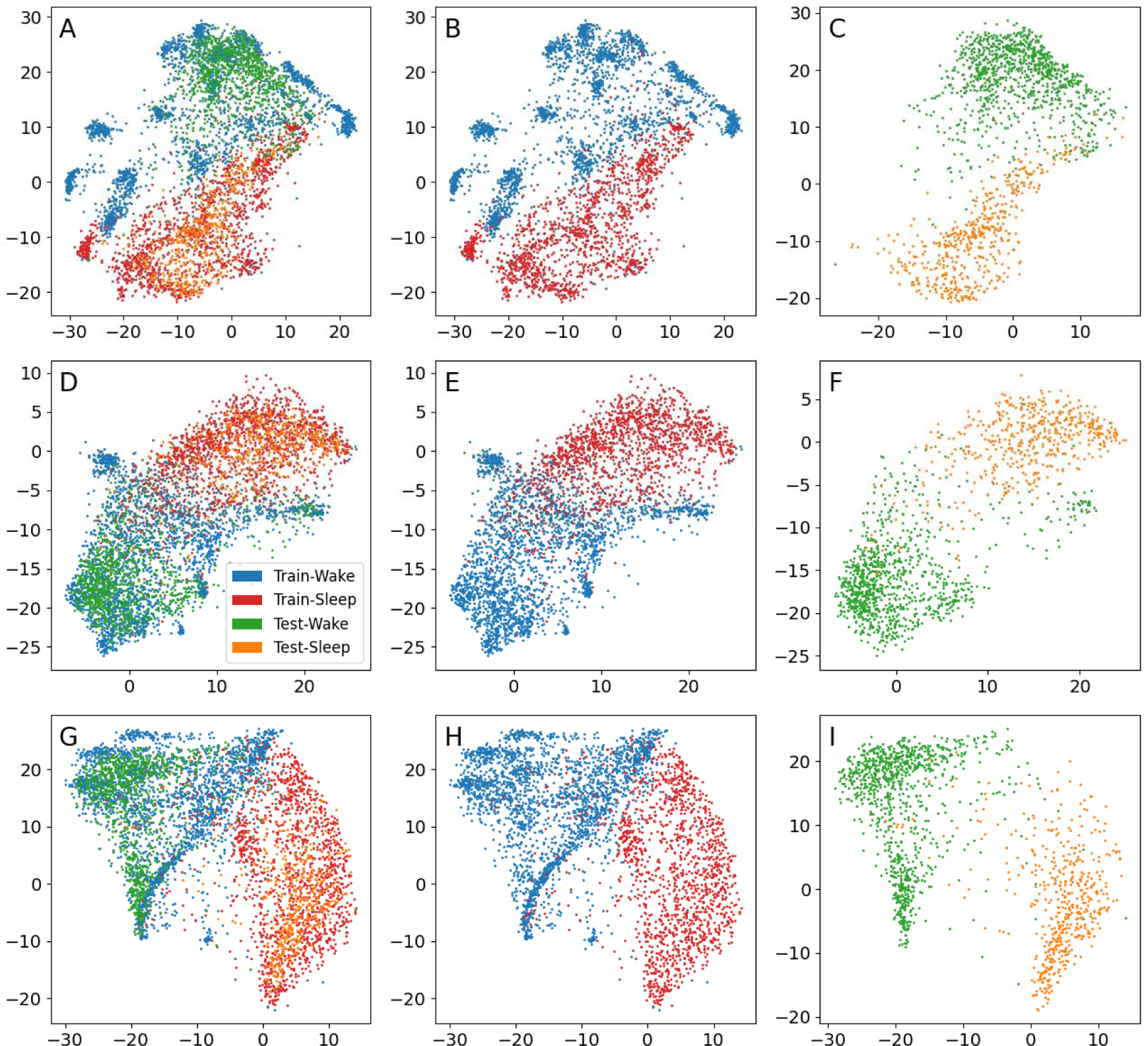
Low dimensional representations for the sleep-wake dataset. Each dot represents 1 s of data. First row (A–C): CNN. Second row (D–F): STFT. Third row (G–I): CWT. First column (A, D, G): All data. Second column (B, E, H): Training data. Third column (C, F, I): Test data.

### 3.2. IEDs

The CNN encoder produced the highest kappa scores (Tab. 2), which were significantly higher for all but for the test data when using an RBF kernel. The CWT encoder scored higher than the STFT encoder, where the latter scored 0 when using a linear kernel.

**Tab. 2.**
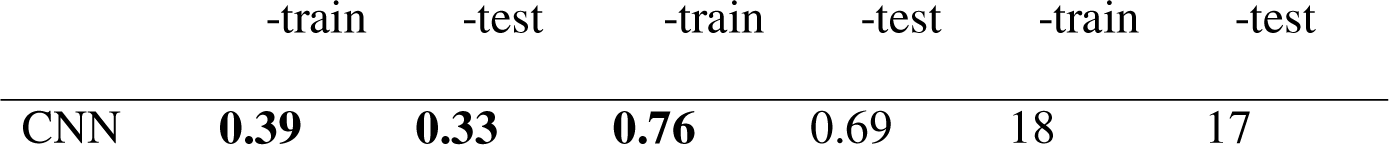
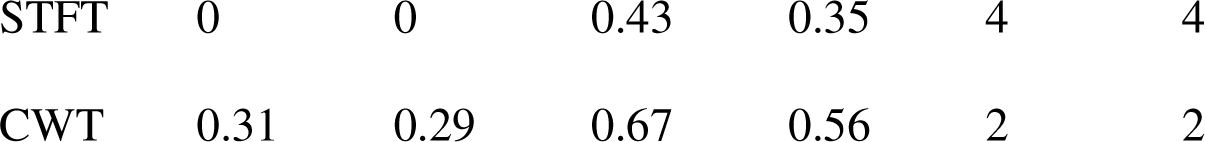
Quantitative measures for the IED dataset. Significantly highest values in a column are in bold (chi-square test). are kappa values for SVM, indicates a linear kernel and a radialis basis function kernel; indicates the number of clusters by k-means.

The CNN encoder produced the highest clustering score for both the training and the test data (Tab. 2), and there were many visually distinct identifiable clusters (Fig. 6B and 6C). Visually, there was some substructure for the STFT (Fig. 6E and 6F) and CWT encoder (Fig. 6H and 6I), but with a tendency for more confluent patterns and mixing of the categories, and the clustering scores were low.

**Fig. 6.**
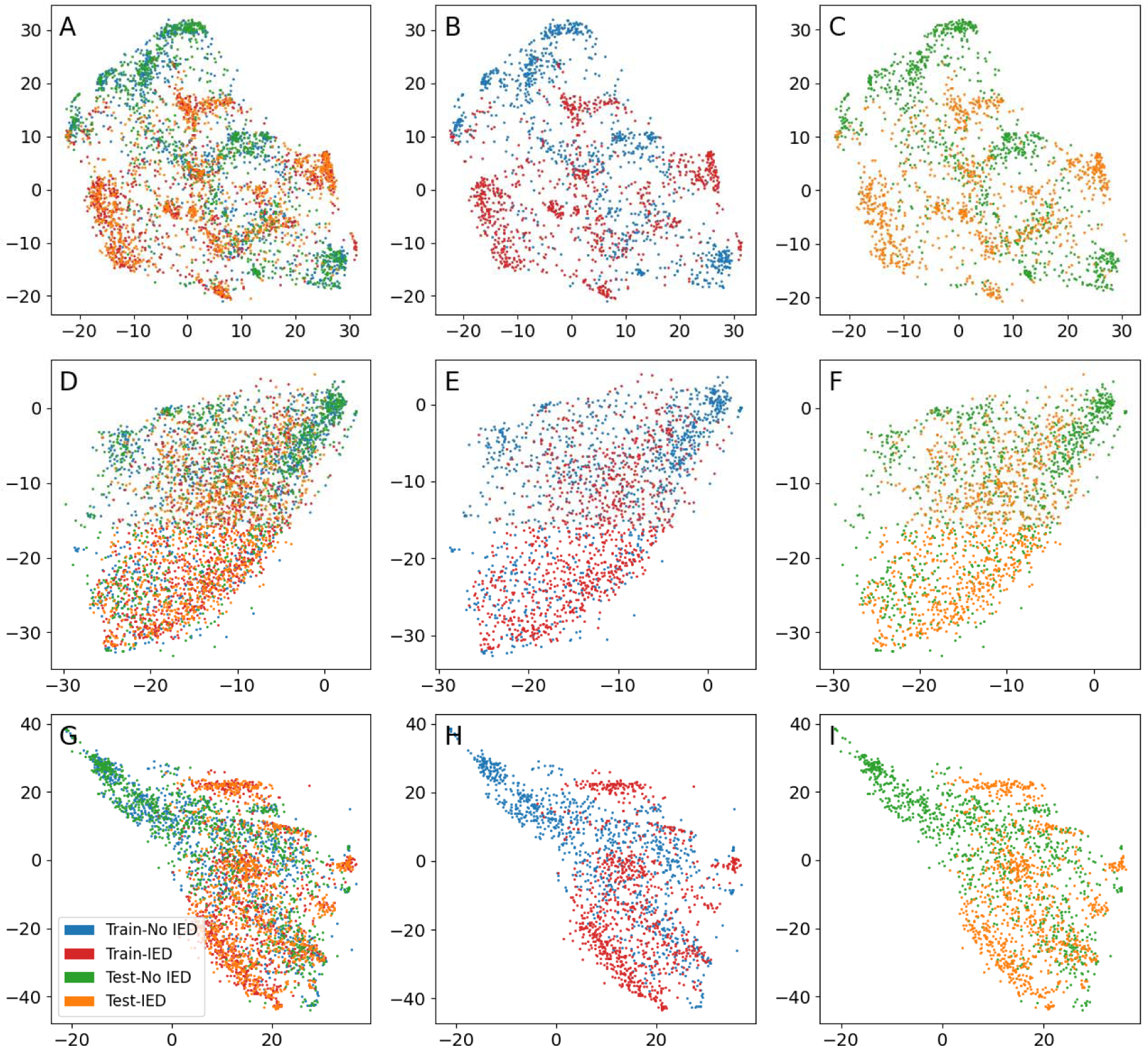
Low-dimensional representation for the IED dataset. Each dot represents 1 s of data. First row (A–C): CNN. Second row (D–F): STFT. Third row (G–I): CWT. First column (A, D, G): All data. Second column (B, E, H): Training data. Third column (C, F, I): Test data.

### 3.3. Seizure activity

Both the CNN and STFT encoder scored 0 when using a linear kernel (Tab. 3). For the RBF kernel, kappa scores were relatively high in general. The CNN encoder scored significantly higher for training using an RBF kernel. There was no significant difference in performance for the methods for test data using an RBF kernel.

**Tab. 3.**
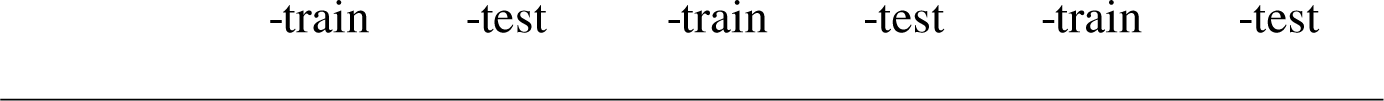
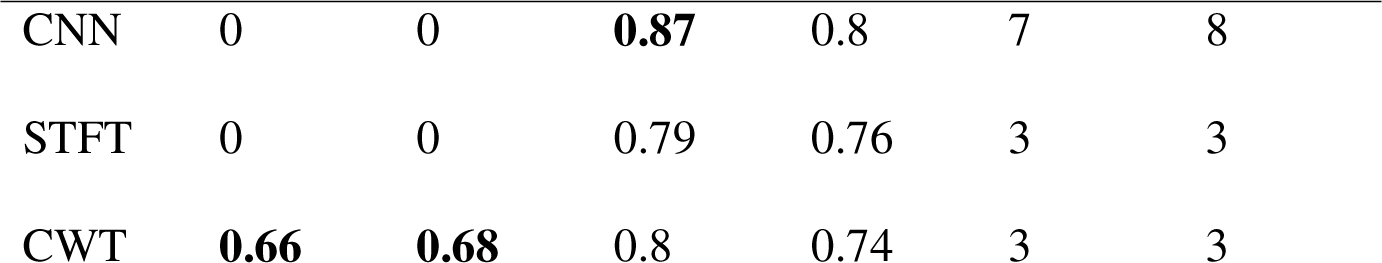
Quantitative measures for the seizure dataset. Significantly highest values in a column are in bold (chi-square test). are kappa values for SVM, indicates a linear kernel and a radialis basis function kernel; indicates the number of clusters by k-means.

|  |  |  |  |  |  |  |
| --- | --- | --- | --- | --- | --- | --- |
| CNN | 0 | 0 | **0.87** | 0.8 | 7 | 8 |
| STFT | 0 | 0 | 0.79 | 0.76 | 3 | 3 |
| CWT | **0.66** | **0.68** | 0.8 | 0.74 | 3 | 3 |

Both with respect to the clustering score (Tab. 3) and the visual appearance (Fig. 7) the CNN encoder performed better for this dataset for both training and test data. The STFT and CWT encoder performed equally well. Visually there were substructures, but the clusters were not distinct enough to produce high clustering scores.

**Fig. 7.**
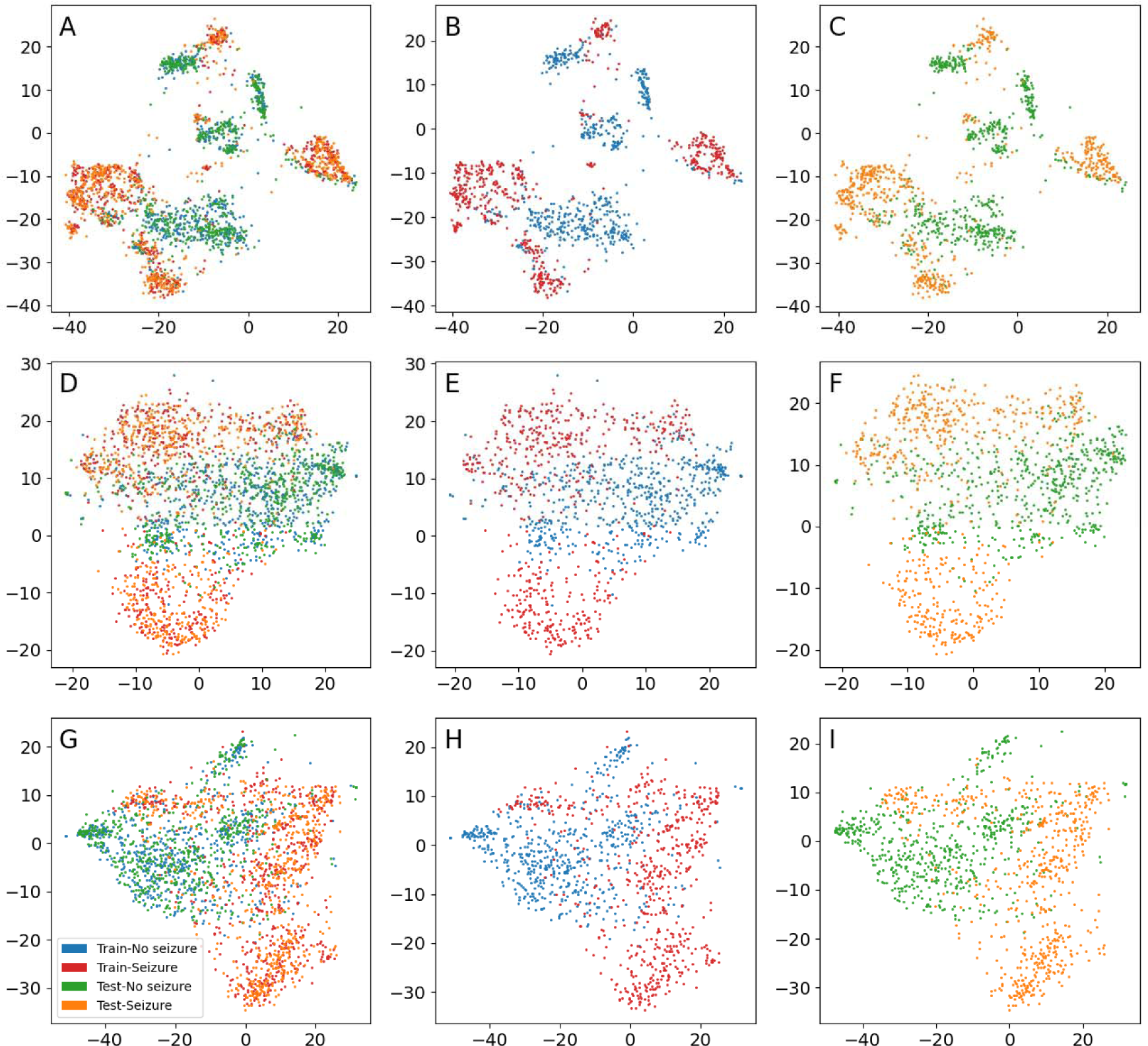
Low dimensional representations for the seizure dataset. Each dot represents 1 s of data. First row (A–C): CNN. Second row (D–F): STFT. Third row (G–I): CWT. First column (A, D, G): All data. Second column (B, E, H): Training data. Third column (C, F, I): Test data.

## 4. Discussion

Convolutional neural networks produced both high- and low-dimensional representations of EEG data and the results were compared to parametric t-SNE using time-frequency features. All methods showed promising results. The most apparent result in favor of the CNN approach was that it produced more distinct clusters of the data in general.

All methods can probably be improved further, for example by optimizing the preprocessing of the data and the hyperparameters of the algorithms. Improvements can likely also be accomplished in a specific sense by tailoring them to each specific dataset and categorization problem, e.g., choice of EEG-channels and frequency bands to analyze. There are different ways of implementing the time-frequency methods, and of course many other methods to consider for producing features when using the original parametric t-SNE. However, given the extent of possible variation of deep learning, it is speculated that this approach has the largest potential for improving the performance.

There was a possible tendency for the suggested approach to perform better for IEDs. As IEDs are short transients, it is possible that the convolutional and the max pooling operations were more efficient in detecting these compared to the CWT and the STFT representations. The CWT encoder seemed to perform better than the STFT encoder for IEDs. The CWT representation had more dimensions than the STFT representation, but it also consisted of statistical values from the wavelet processing, where the max value was equivalent to the max pooling operation used in the level analysis of the CNN encoders, and this may have contributed to it performing better for IEDs than the STFT representation.

As evaluated by the SVMs and kappa scores, the methods showed similar performance for the sleep-wake and the seizure dataset. These categories are to a large extent defined by their frequency content, making it reasonable that time-frequency methods can perform well for these data types.

In contrast to the IED and seizure datasets, the number of clusters decreased for the test data compared to the training data of the sleep-wake dataset. The main difference between the datasets, apart from the difference in EEG patterns, was that the test data of the sleep-wake dataset consisted of new subjects, whereas for the other two datasets, the test data consisted of new data from the same subjects that were used for training. This implies overfitting to the training subjects, however, the global structure generalized to new subjects. Further testing to elaborate if the higher clustering ability of the CNN encoders were due to overfitting to the subjects is needed. Such testing may involve using overlapping examples when training which, as demonstrated in Appendices A.3 and E, can have an impact on the results. Training and testing on larger datasets are also necessary.

If there is overfitting, it is probably based on the distance between the signals. This is suggested by all low-dimensional representations produced by the method in this article since the resulting clusters conform with the annotations. Hence, the low-dimensional representations of the training data can still be useful and give insight into the relations between those specific signals even when the encoders do not generalize well to new data. Whether it is worth the time and resources to train unique encoders to every dataset as a method of analysis is another question.

The suggested approach can be time consuming to develop. If a high-dimensional representation that produces good results for t-SNE can be identified, then there are faster implementations for the algorithm, e.g., Barnes-Hut t-SNE (van der Maaten, 2014) and Flt-SNE (Linderman et al., 2019), and the deep learning approach may be unnecessarily complicated.

As it is, some knowledge in programming is necessary to implement the method; even using ready-made scripts from this work will require some understanding of Python and Tensorflow, and willingness to invest time to understand and set up the workflow and solve potential problems with mismatching software dependencies.

Since the method is self-supervised, anything in the data may induce clustering. This may be beneficial, e.g., the method will in a sense be more unbiased. On the other hand, it may also make the method too unselective, e.g., uninteresting artifacts may cause clustering of data that is suboptimal for the task at hand. The method could also be too sensitive, e.g., theoretically, small variations in the electrode placement could affect the generalization if training and test data are recorded at separate occasions; or there may be spurious clustering induced by noise.

As described in the introduction, t-SNE has previously mainly been used to visualize EEG data. There are several promising applications: The method could be further developed to produce quick overviews of whole recordings, providing a rapid first assessment evaluating if an EEG is normal or pathological. This could be useful to, e.g., prioritize the order of the assessment of EEGs in the clinical routine work when workload is high. The method might also be used to construct trends for long term monitoring. For example, seizure activity and status epilepticus could easily be detected in intensive care units lacking personnel with training in EEG interpretation. Jing et al. (2019) integrate visualization using t-SNE in an annotation tool; rapid annotation of data could be performed by simply manually marking clusters and assign categories in a graphical user interface, which then automatically annotate the corresponding time intervals in the EEG.

## 5. Conclusions

Using CNNs the data tend to group or cluster in agreement with the traditional EEG categories which is a first step towards further improvements of EEG categorization. The CNN method shows potential for use in several clinical areas, in addition to research.

## Acknowledgments

The work was mainly funded by Linköping University, the University Hospital of Linköping, and the ALF of Region Östergötland (LIO-936176 and RÖ-941359). AE was supported by the ITEA3/VINNOVA funded project (ASSIST). We would also like to thank the Nvidia corporation, who donated the Quadro P6000 and Titan X graphics cards used for training the encoders.

## Author contributions

Mats Svantesson: Conceptualization; Formal analysis, Investigation, Methodology, Project administration, Resources, Software, Visualization; Writing - original draft; Writing – review & editing. Håkan Olausson: Writing - review & editing. Anders Eklund: Resources, Supervision; Writing - review & editing. Magnus Thordstein: Writing - review & editing.

## Appendix A. Data

### A.1. Data selection and annotation

Three datasets for clinically significant EEG activities were created to test the proposed method: 1) sleep vs. wakefulness, 2) IEDs vs. no IEDs, and 3) seizure activity vs. seizure-free activity.

To produce the project’s database, a set of 1,000 EEGs from 1,000 unique subjects were first extracted from the TUH EEG Corpus (Obeid and Picone, 2016) by automated search. Inclusion criteria were subject age ≥18 years, sampling rate 250 Hz, and duration of at least ≥ 600 s. This resulted in a set of subjects with an average age of 51 years (553 females) and recordings with an average duration of 1356 s. From this database, a subset of EEGs containing: 1) wakefulness, 2) sleep, 3) frequent IEDs, and 4) seizure activity was selected. The number of EEGs was limited by their natural occurrence in the extracted material. For category 2, the EEGs were dominated by sleep, but it was difficult to find complete EEGs without some amount of wakefulness. These EEGs were therefore dichotomously annotated as 10 s epochs so that each epoch represented wakefulness or sleep. Sleep was defined according to the American Academy of Sleep Medicine’s (AASM) sleep scoring manual (Iber et al., 2007) but modified to 10 s instead of 30 s and sleep stages were not indicated. IEDs were annotated according to criteria proposed by Kane et al. (2017). Seizure activity was defined according to Hirsch et al. (2021). Selection and annotation were performed by a specialist in clinical neurophysiology (author MS).

The above procedure resulted in 19 EEGs of wakefulness, 14 EEGs of dominating sleep, 31 EEGs with frequent epileptiform activity, and 6 EEGs containing seizure activity. From these EEGs, the datasets were created with examples of 1 s duration and the sets where split with probabilities 0.75 for training and 0.25 for testing respectively. The sleep-wake dataset had the largest number of subjects and examples. This dataset was split with respect to subjects, i.e., testing would evaluate generalization to new subjects, which resulted in 14 awake and 5 sleeping subjects for training and 8 awake and 3 sleeping subjects for testing. The IED and seizure datasets were split with respect to examples, i.e., testing would evaluate generalization to new data from the same subjects. A summary of the number of examples per dataset and the respective categories of the datasets is given in Tab A.1.

**Tab. A.1.**
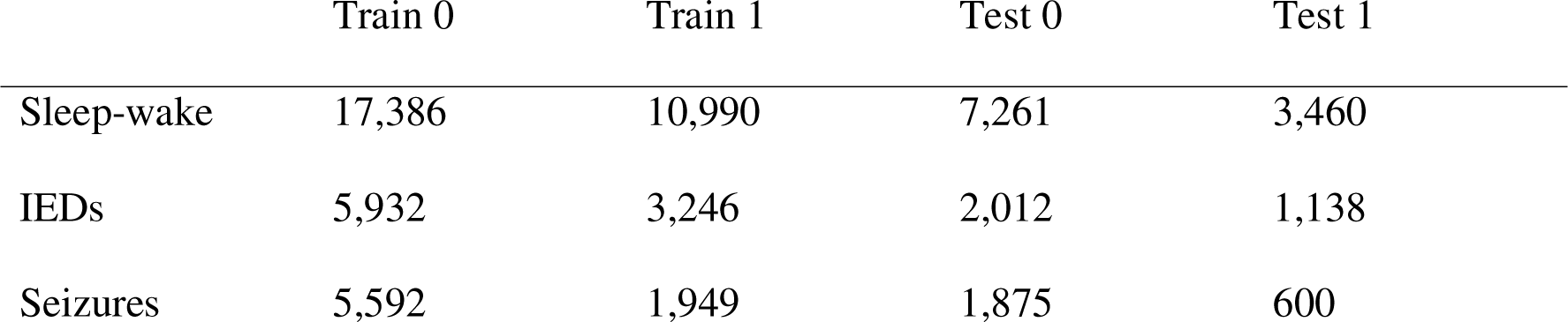
Summary of the number of examples per dataset and category using examples of 1 s duration. Sleep-wake: 0 = wake, 1 = sleep. IEDs: 0 = no IED, 1 = IED. Seizure: 0 = no seizure activity, 1 = seizure activity.

|  | Train 0 | Train 1 | Test 0 | Test 1 |
| --- | --- | --- | --- | --- |
| Sleep-wake | 17,386 | 10,990 | 7,261 | 3,460 |
| IEDs | 5,932 | 3,246 | 2,012 | 1,138 |
| Seizures | 5,592 | 1,949 | 1,875 | 600 |

These datasets had not been used in the previous development of the method. For this, a few of the 1,000 extracted EEGs, not used as training or test data for the results, were used for preliminary testing of the Python scripts before the final analysis using the three datasets. Especially one longer recording was utilized and it was also used for the demonstration in Appendix E. One of the seizure EEGs, representing status epilepticus, was reused here in the appendices to generate the illustrative examples.

### A.2. Example duration

To be able to identify patterns, it is necessary to have a sufficient number of data points, as is the case when training deep learning models. For a standard 20-minute recording, an example duration of 1s give 1,200 unique examples, compared to, e.g., 40 unique examples when using an example duration of 30 s. For instance, if an EEG contains one 15 s episode of seizure activity, an example duration of 1 s will give 15 examples, whereas a longer duration may only give a few examples or that the activity may only constitutes part of the duration of an example. For EEGs with frequent IEDs, a longer duration could result in almost all examples containing IEDs making it hard to study this waveform. Most waveforms of interest will fit within 1 s but some patterns, e.g., seizure activity and sleep stages, may become poorly represented. For sleep stages, during an annotated epoch, the character of the activity can change so that different parts of the epoch could represent different sleep stages. Thus, using 1 s examples with data annotated for 10 or 30 s epochs probably results in some examples being mislabeled. However, annotating sleep for 1 s epochs would be very time consuming and probably produce a large scoring variability since there is no clear definition for sleep staging of such short time intervals. The chosen duration of 1 s was thus a compromise, mainly to allow for a larger number of examples for training and testing. For the long duration patterns, some degree of label noise was expected. A demonstration of using a longer duration is given in Appendix E.

### A.3. Overlapping vs. non-overlapping examples

A way to counteract overfitting in machine learning is to use overlapping examples, thereby increasing the variation of the training data. Shorter signal transients, e.g., IEDs, may appear at any time position in an EEG. Therefore, using a limited amount of non-overlapping examples may result in encoders only detecting transients at certain positions in an example. The effect on the result of using overlapping vs. non-ovelapping examples may sometimes be pronounced (Fig. A.2). The effect on generalization is demonstrated in Appendix E.

**Fig. A.2.**
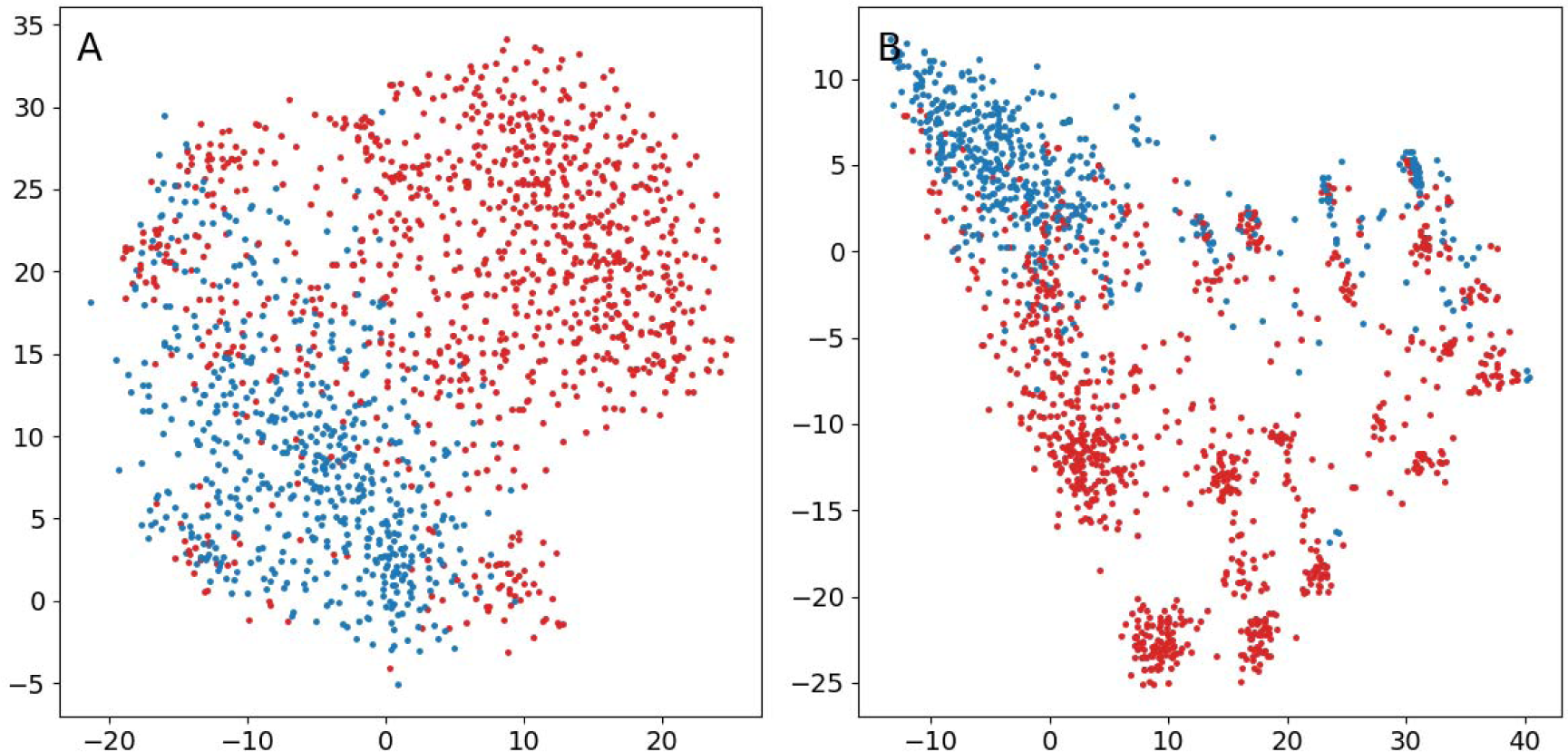
Comparison of training encoders using non-overlapping or overlapping time intervals as examples. EEG of 1,601 s duration representing status epilepticus where each dot represents 1s of data and red dots indicate seizure activity. (A) Non-overlapping examples. The EEG was divided into 1,601 unique examples by using consecutive 1 s intervals. (B) Overlapping examples. The whole EEG-recording was loaded into CPU memory, examples were then generated online during training by randomly selecting time positions in the recording and extracting 1 s of data.

As described below (D.2), time-frequency methods were used as a comparison in the evaluation of the method. Here, using overlapping examples, the amount of data can expand substantially. As an example, a 21-channel 20 min EEG-recording with sampling frequency 250 Hz has 6,300,000 data points. If all data are used to create non-overlapping examples, then this will give 1,200 unique examples. The total data size will be the same if the whole recording is used to generate random overlapping examples online during training. A time-frequency method that represents each example by 630 values will reduce the amount of data when using non-overlapping unique examples (756,000 data points) but increase it if using overlapping examples and keeping the time resolution (189,000,000 data points). Using randomly selected overlapping examples therefore proved hard to implement for the time-frequency methods. For the present study, fixed non-overlapping examples were used in all instances to make comparisons more fair. This limitation of the study was thus due to the time-frequency methods utilized.

## Appendix B. t-SNE

### B.1. The original t-SNE

In the original t-SNE developed by van der Maaten and Hinton (2008), the joint probability *p_ij_* of a pair of high-dimensional data points *x_i_* and *x_j_* being neighbors is assumed to follow a normal distribution according to

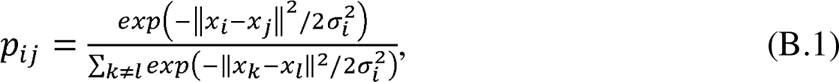

where 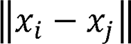 is the Euclidean distance *d_ij_* and *σ_i_*is the standard deviation. *σ_j_* is determined by a binary iterative search to meet a condition imposed by the *perplexity* parameter. The perplexity is 2*H*(*P_i_*), where *P_i_* is the probability distribution corresponding to *σ_i_* and 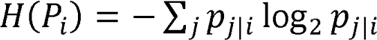 is the Shannon entropy. Thus, the perplexity will determine *σ_i_* and reflects the number of efficient neighbors. In the original implementation, the diagonal is set to zero (*p_ij_* = 0, *i* =*j*) and the symmetrical matrix 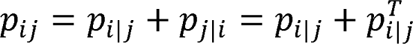 is used.

For calculating the probability *q_ij_* of a corresponding pair of low-dimensional data points *y_i_*and *y_j_* being neighbors, a t-distribution was introduced by van der Maaten and Hinton (2008) as

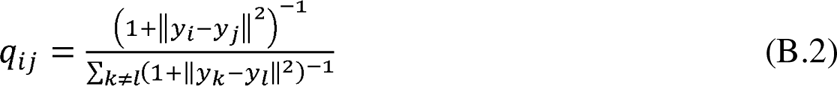

To match the resulting probability distributions b and 6, the Kullback-Leibler divergence

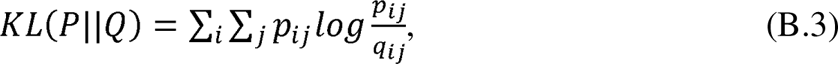

which reflects the difference between the distributions of the probabilities, is used as loss function. Gradient descent is used to minimize the loss function.

### B.2. Modified t-SNE using custom probability distributions

The difference between the normal distribution and the t-distribution, i.e., that the t-distribution has heavier tails, is supposed to create favorable attractions and repulsions, resulting in preservation of local structure while overcoming the crowding effect (van der Maaten and Hinton, 2008) (Fig. B.1A). In the suggested approach of the present work, the high-dimensional representation was generated by the encoder. The representation thus changed during training and the corresponding probabilities had to be recalculated for each training batch. The original t-SNE uses an iterative search to determine the standard deviation of the normal distribution given the perplexity. This can be time consuming (see Appendix A.6), and a custom distribution based on ranked distances was therefore used instead.

**Fig. B.1.**
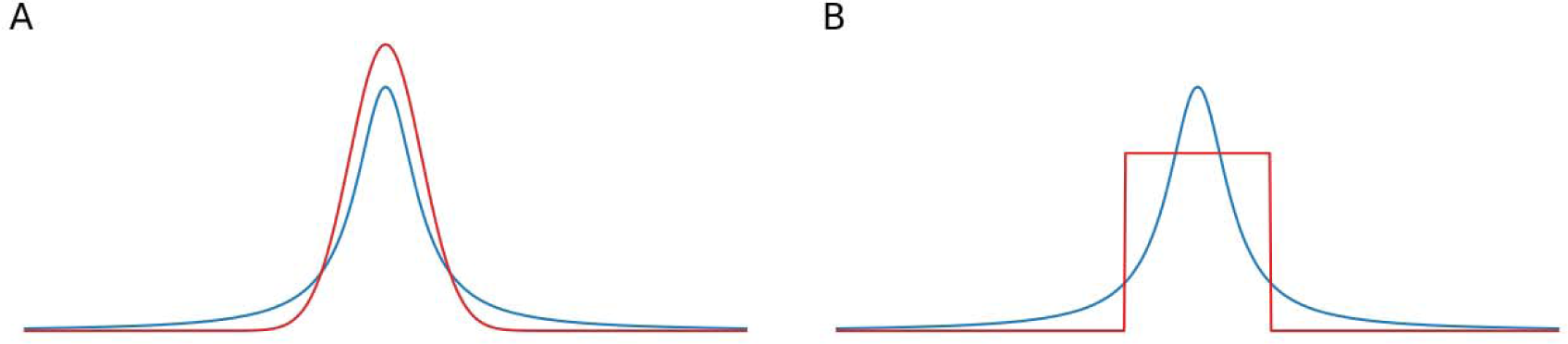
Theoretical probability distributions. The blue curve is the t-distribution with one degree of freedom. (A) The normal distribution in red. (B) The suggested distribution in red.

By ranking the distances between examples, custom distributions based on the rank can be constructed. A simple alternative is to assume that each example has a certain number of close neighbors, and that all other examples are distant. Thus, by setting the lowest ranked distances to a value, the rest to a value, then normalizing with the sum of all values this resulted in a probability distribution with probabilities (Fig. B.1B.)

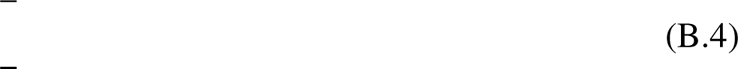

If the number of examples is, then. In this approach, the information regarding distances and ranks is lost.

### B.3. Comparison of using a normal or a distribution based on ranked distances in the original t-SNE

To illustrate the performance using the suggested distribution, the original t-SNE was performed using either its original implementation with a normal distribution or modified for using the suggested distribution. The code of van der Maaten for a Python implementation of t-SNE was used (https://lvdmaaten.github.io/tsne/code/tsne_python.zip) to perform the tests. A modified version was created for the suggested distribution. The original code has a default value of 50 components for PCA, which was used. The low-dimensional representation was initialized randomly from a uniform distribution and the same randomization was used in all examples. Early exaggeration was used with a factor of 12 for the first 250 iterations, as suggested by Kobak and Berens (2019), and 5,000 iterations were performed. t-SNE was performed for an EEG using a time-frequency representation (STFT; Appendix D.2.1) for perplexities 0.25, 1, 5, and 20 percent.

The resulting low-dimensional representations were similar (Fig. B.2). For higher perplexity values the suggested distribution seemed to produce less structure, i.e., more confluent patterns. However, it is possible that the perplexity and the number of neighbors are not equivalent. The range of values for the low-dimensional representations differ, where the range decreases more rapidly with increasing number of neighbors for the suggested distribution compared to the normal distribution.

**Fig. B.2.**
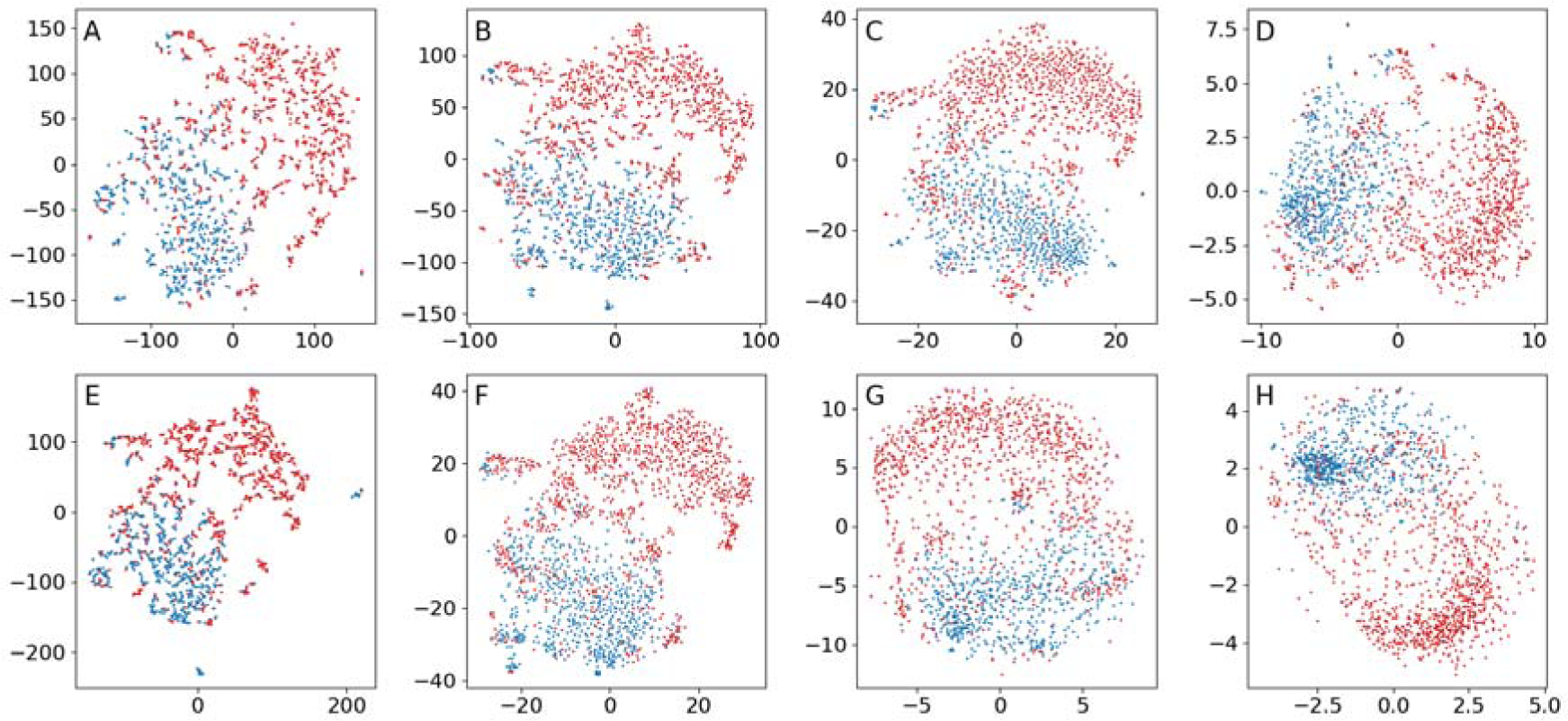
Comparison of the original t-SNE performed with the suggested or a normal distribution. The data is a short-time Fourier transform representation of an EEG representing status epilepticus where each dot is 1 s and red dots indicate seizure activity. First row (A–D): using a normal distribution. Second row (E–H): using the suggested distribution. (A): perplexity 0.25 %. (B): perplexity 1 %. (C): perplexity 5 %. (D): perplexity 20 %. (E): 0.25 % neighbors. (F): 1 % neighbors. (G): 5 % neighbors. (H): 20 % neighbors.

### B.4. Comparison of using a normal or a distribution based on ranked distances when training encoders

To illustrate the effects on performance of using a normal or the suggested distribution when training encoders, parametric t-SNE encoders (Appendix D2) and CNN encoders (Appendix C) were trained using either of these distributions. A batch size of 500 was used and perplexity and the number of neighbors were both set to 5. All resulting low-dimensional representations showed a strong division according to the categories, but possibly a tendency for more local structure when using the suggested distribution (Fig. B.3).

**Fig. B.3.**
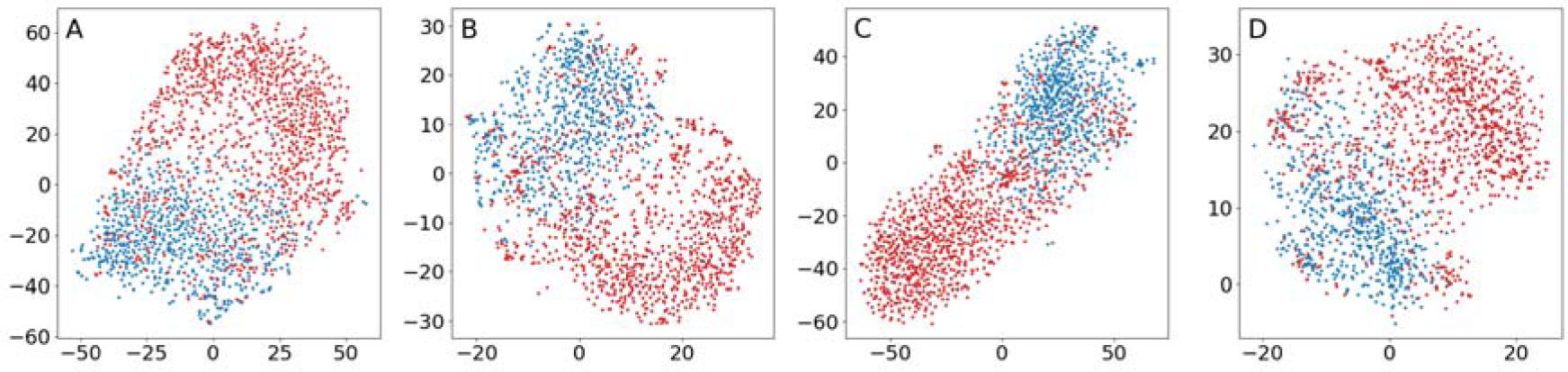
Comparison of parametric t-SNE and CNN encoders when using the normal and the suggested distribution. Batch size during training 500, perplexity/neighbors of 5. (A) Parametric t-SNE with a normal distribution and STFT features. (B) Parametric t-SNE with the suggested distribution and STFT features. (C) CNN encoder with a normal distribution. (D) CNN encoder with the suggested distribution.

Generally, best results were produced using low values for the number of neighbors u (0.5–2 percent of the batch size) (Fig. B.2). In the article, u = 5 was used, which for a batch size of 500 is equal to 1 percent. This is in line with the rule of thumb for the value of the perplexity suggested by Kobak and Berens (2019).

### B.6. Time consumption for a normal and a distribution based on ranked distances

The algorithms for estimating the normal and the suggested distributions were performed for examples having 50 components and batches of 500, 1,000, 2,500, 5,000, 10,000, 20,000, and 50,000 examples. The tests were performed on a computer with an Intel Core i7-6800K CPU 3.4 GHz (6 cores/12 threads).

In computing the suggested distribution, the distance involves matrix multiplication (*O*(*n*^3^)), and the sorting on average should take O((*n*log*n*)) but in the worst case may take *O*(*n*^2^) (https://numpy.org/doc/stable/reference/generated/numpy.sort.html) and there are *n* instances of sorting. In the tests, the processing times using either distribution followed the same time complexity but using a normal distribution consumed two orders of magnitude more time (Fig. B.4). However, since computing the distributions only constitutes part of all computations during training, for a batch size of 500 training times were 2–2.5 times faster when using the suggested distribution.

**Fig. B.4.**
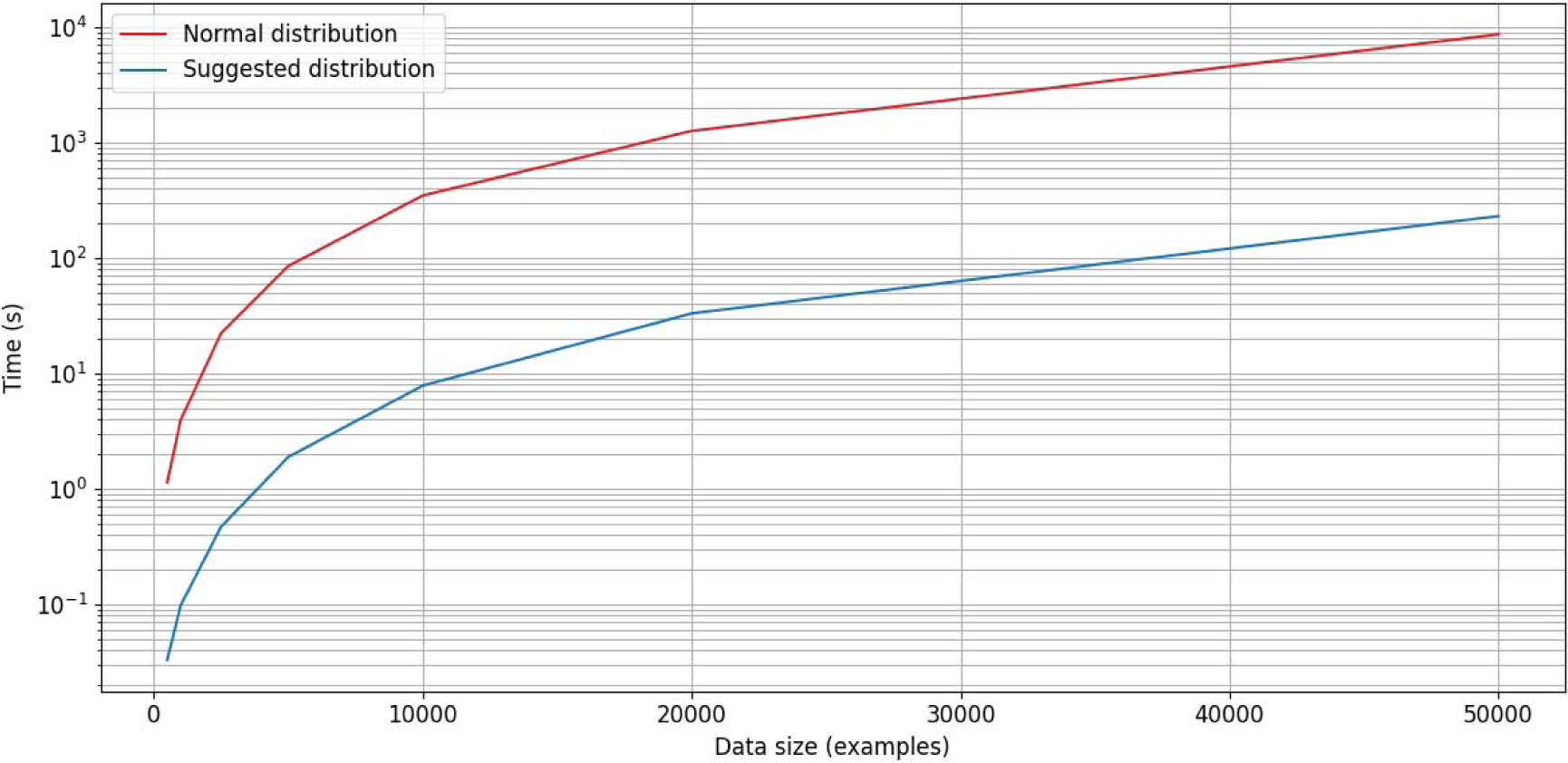
Comparison of processing times for estimating the normal distribution (red) and the suggested distribution (blue) in the present work. Estimations are 34–47 times faster when using the suggested distribution. However, in practice, the estimation only constitutes part of the computations during training, the training time will depend on the batch size, and for a batch size of 500 training times was often approximately 2–2.5 times faster when using the distribution.

## Appendix C. Encoders

### C.1. Encoder architecture

In all encoders the input was processed through a series of blocks of convolutions and max pooling with the intention of producing levels with increasing field of view of the input (Fig. C.1). Each level was then analyzed by further convolution, max pooling followed by fully connected layers to generate a set of values characterizing it. The values from all levels were concatenated and became the latent representation. Each EEG channel was analyzed separately throughout the convolutional blocks and level analysis. A fully connected block of four layers then mapped to a two-dimensional representation . To be able to access the latent representation, all types of encoders were implemented in two parts–from to and from to . Putting the two parts in series produced the full encoder.

**Fig. C.1.**
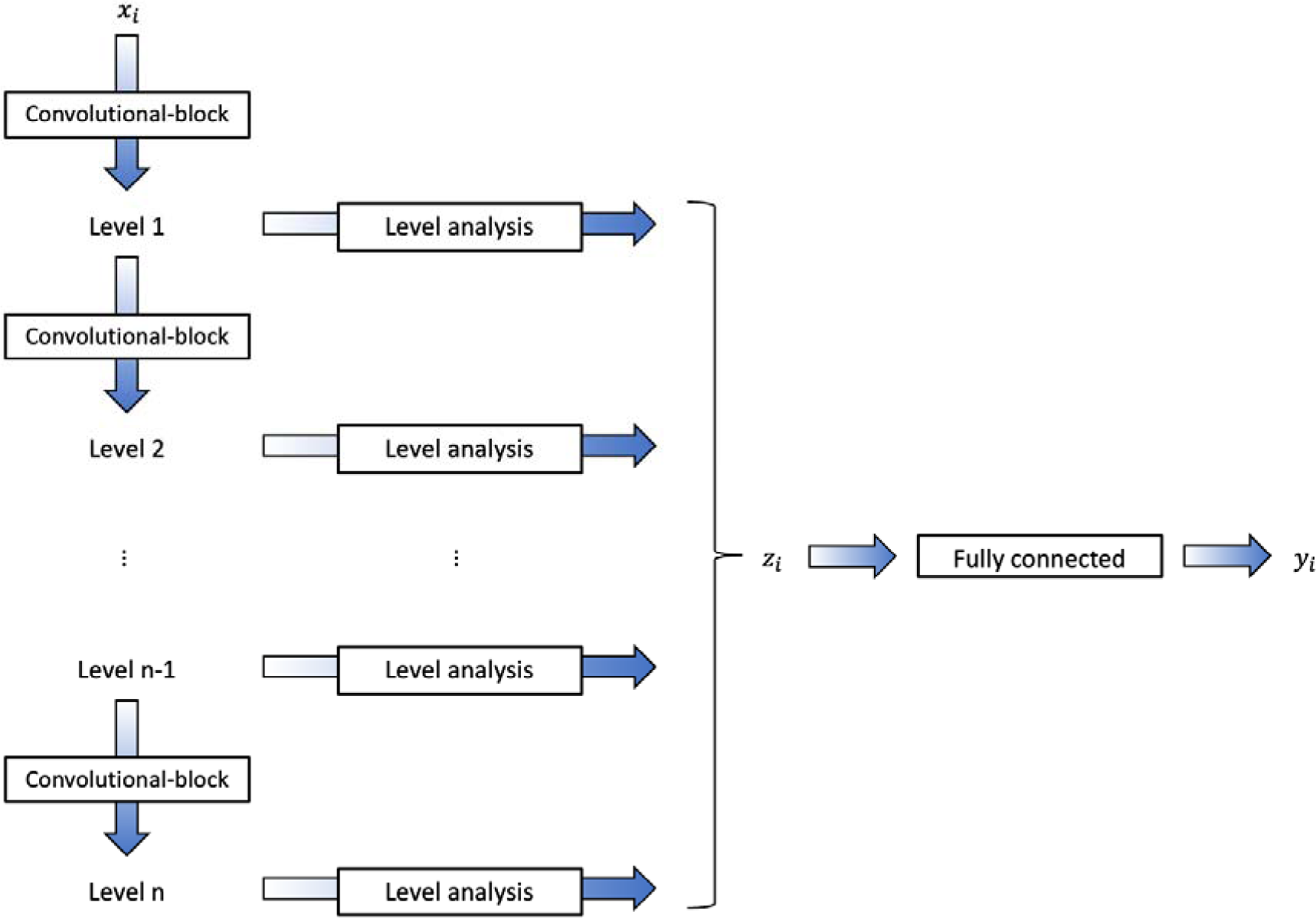
Overview of the encoder architecture. To the left, a series of convolutional blocks processed and successively downsampled the data through max pooling to produce levels of increasing field of view. Each resulting level was analyzed further and the results from all levels were concatenated to produce a latent representation of the data . A series of fully connected layers then produced the low-dimensional representation .

#### C.1.1. Convolutional blocks

The convolutional blocks (Fig. C.2A) consisted of three convolutional layers, each with 16 filters and strides of 1. The first layer had a kernel size of 2, the second and third kernel sizes of 1. Scaled exponential linear units (SELU) (Klambauer et al., 2017), which have self-normalizing properties, were used as activations between the layers. Each block was finished with max pooling with a kernel size of 2 and strides of 2, thus downsampling by a factor of 2.

**Fig. C.2.**
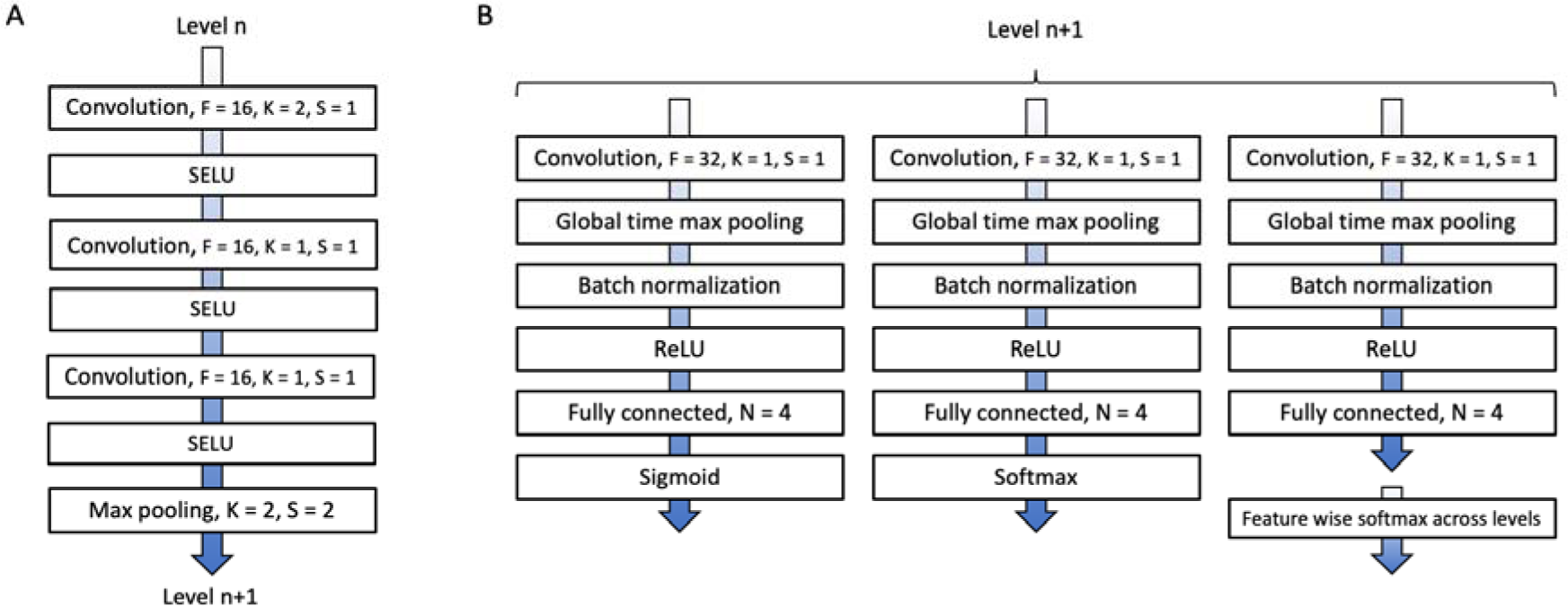
(A) Illustration of a convolutional block. (B) Illustration of the level analysis. F: number of filters; K: kernel size; S: strides; N: number of nodes. ReLU: rectifying linear unit.

#### C.1.2. Level analysis

One aspect of the idea of the encoder architecture was that the resulting values from the level analysis would represent the presence of high-level waveform features at different field of view. It was assumed that some of these could be independent features, but there could also be dependent features, either being mutually exclusive or of different importance. Dependence could be either for a certain level or across different levels. For example, from the EEG experts perspective, there may be artifacts but at the same time visible cortical activity, activity can be classified as either regular or irregular, and waveforms with pointy morphology may be of greater importance than those with round morphology.

As an implementation of this idea, the level analysis consisted of three parallel blocks (Fig. C.2B), each starting with a convolutional layer (32 filters, a kernel size of 1, and strides 1), followed by max pooling across the time dimension reducing this dimension to 1, then batch normalization, a rectifying linear unit activation, and a fully connected layer (4 nodes). The first block ended in a sigmoid activation, and this would constitute independent features (Fig. C.3, left). The second ended in a softmax activation, and this would constitute intra-level dependent features. (Fig. C.3, middle) From the third block of all levels, groups were formed consisting of one feature from each level, and each group were put through a softmax activation, and this would constitute inter-level dependent features (Fig. C.3, right).

**Fig. C.3.**
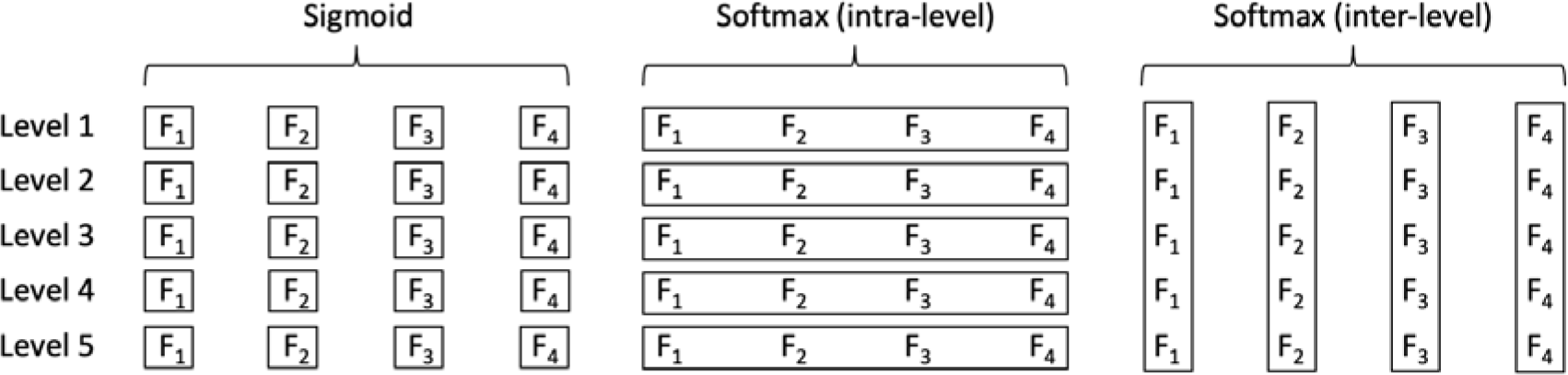
Illustration of the activations for the latent representation for 5 levels. Boxes indicate dependence; for the sigmoid activations, each feature F_i_ can take any value between 0 and 1 independent of other features; for softmax activations, the values of the features in each box are normalized to one so that

For downsampling by a factor of 2 and the sampling frequency of 250 Hz of the original data, a time step of the first and second level would correspond to 8 ms (125 Hz) and 16 ms (62.5 Hz), respectively. Since the idea was to create an encoder that mimic how the expert visually analyzes an EEG, the data was low pass filtered at 40 Hz (which is used traditionally by most clinical neurophysiologists at the authors’ clinic), the first two levels were skipped, and the remaining 5 levels (32, 64, 128, 256, and 512 ms) were used. This resulted in a latent representation of 1260 dimensions.

#### C.1.3. Fully connected block

The mapping of to was accomplished using three fully connected layers, each with 512 nodes, followed by a final fully connected layer with the 2 nodes (Fig. C.4). SELU activations were used in between the fully connected layers.

**Fig. C.4.**
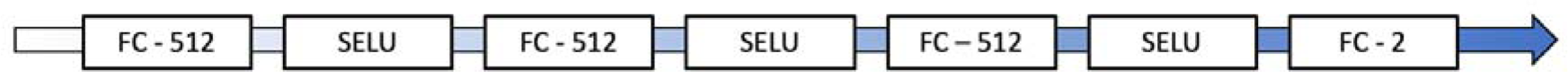
Illustration of the fully connected block mapping to . FC: fully connected layer with the number of nodes indicated.

### C.2. Loss function

Before training on a batch, the high-dimensional representation had to be predicted by the first part of the encoder (Fig. C.5). From this, the probability distribution could be computed.

**Fig. C.5.**
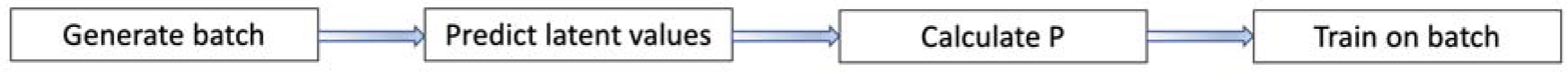
Illustration of the pipeline for generating a training batch. First, examples were randomly selected. The latent values were then predicted using the first part of the encoder. Thereafter, the probability distribution was calculated, and and were then feed into the full encoder for training.

The loss function thus consisted of calculating the resulting probability distribution of the low-dimensional representation, given by (B.2), and then the Kullback-Leibler divergence of and, given by (B.3).

### C.3. Hyperparameters

The method described in this paper, was developed over a long period of time using various data. The encoder architecture slowly evolved and emerged as an alternative that on average produced consistent and interesting results. When training and using validation data, at least for small datasets, there were usually some degree of overfitting to the training data. However, it was almost always the case that if the training loss converged, then the validation loss would also converge. Given this previous experience, in the present evaluation, no validation data was used during training. Only the training loss was monitored, and the only parameter considered for adjustment was the learning rate.

The number of filters were limited by memory constraints; the extraordinarily small kernels were chosen to reduce the memory load. The number of nodes of the fully connected layers of the level analysis was selected to give a high-dimensional representation with a size in the same range as the comparative methods (630–2,436 dimensions; Appendix D2.1-2). The number of nodes of the fully connected layers of the mapping from *z_i_* to *y_i_* was chosen to have a total number of parameters that was less than for the parametric t-SNE encoders (D.1). These last two choices were made to make comparisons with the parametric t-SNE more fair. The first part of the encoder had 19,084 parameters (18,124 trainable) and the second part had 1,171,970 parameters (all trainable).

Adam was used as optimization algorithm (Kingma and Ba, 2015) with parameters β1 = 0.5 and β2 = 0.9. All convolutional and fully connected layers used L2 kernel regularization (default value of 0.01), and all kernels of these layers that were followed by SELU activations were initialized using the LeCun normal distribution as recommended in the tensorflow documentation (https://www.tensorflow.org/api_docs/python/tf/keras/activations/selu). A batch size of 500 was used.

A learning rate of 1e-4 was used for the data utilized to produce the results and all examples of low-dimensional representations, except for the sleep scoring data when training with overlapping examples in Appendix E, where 5e-6 was used. The loss will show some smaller variation of random character (Fig. C.6A), which is mostly due to variation of the data content from batch to batch. If there are fluctuations of the loss’ baseline with local trends, then the encoder is probably moving between different local minima and training may not satisfactorily converge (Fig. C.6B); if the low-dimensional representation is visualized intermittently during training, then it may show dramatically changing topology. However, the training may still stabilize, producing a satisfactorily result (Fig. C.6C).

**Fig. C.6.**
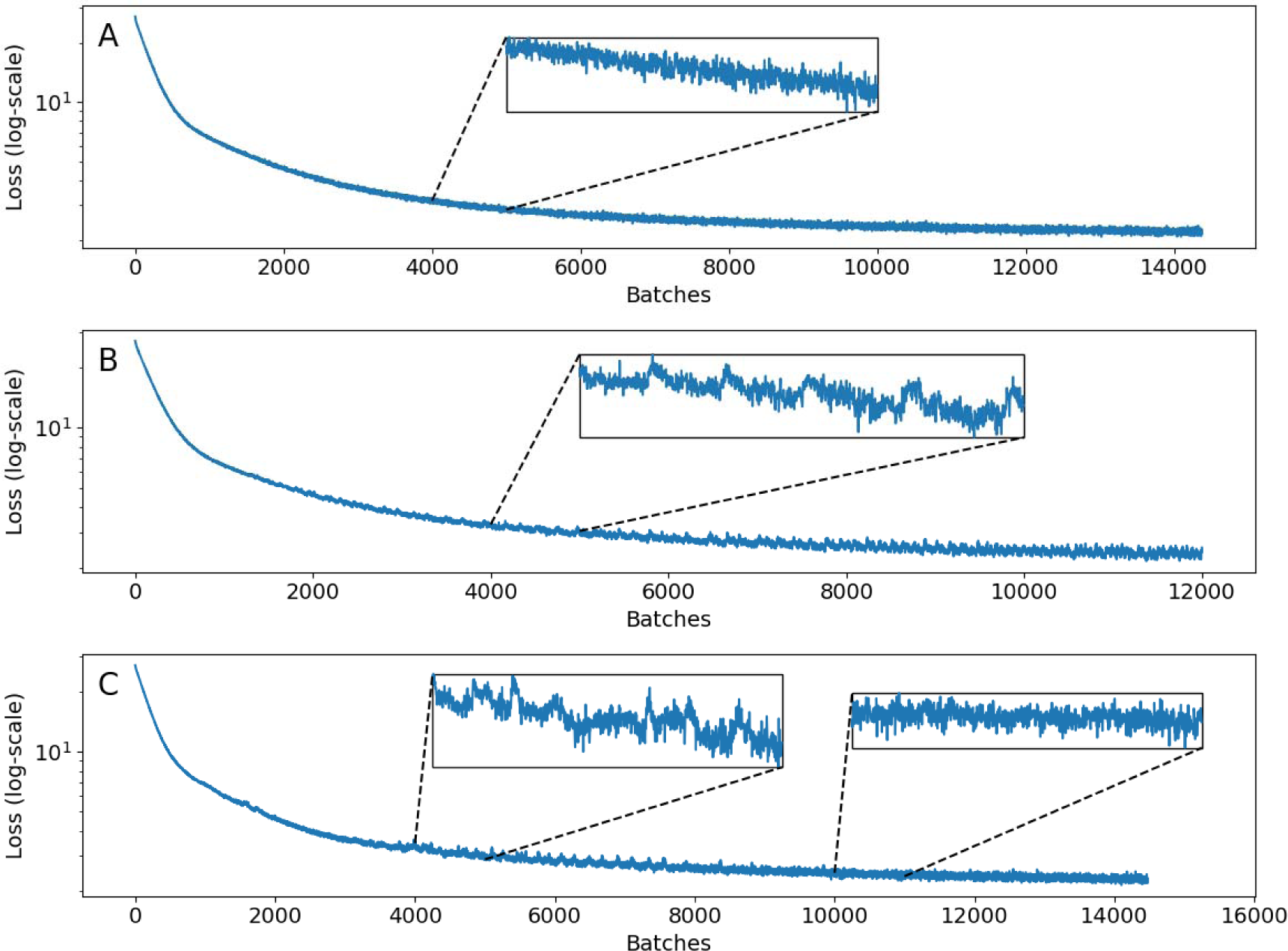
Examples of training losses. (A) Example where the loss decreases steadily apart from smaller fluctuation of random character. (B) Example where the loss shows instability in the form of a low amplitude sawtooth pattern which represents movement between local minima with different topology for the low-dimensional representation. (C) Example where the loss shows transient instability but then stabilizes and converges to one topology.

A general recommendation would be to start training with a learning rate of 1e-4. If the loss does not decrease steadily and converge, then restart using a lower learning rate. Restarting several times using incrementally lower rates (e.g., 5e-5, 1e-5, 5e-6, …) may be necessary.

Training was performed until the loss stopped decreasing. This point was decided by comparing the average loss of the last 2,500 iterations with the preceding 2,500 iterations to a precision of 2 decimals. The optimal length of the intervals over which the averages are calculated that works best will depends on the data and the learning rate. So, the intervals may have to be increased when using a smaller learning rate to prevent premature stopping or decreased for a specific dataset to avoid continue training after the loss has converged satisfactorily. The resulting training times for the three datasets were 12–13 h each when using a single GPU.

## Appendix D. Comparative methods

### D.1. Parametric t-SNE

The original implementation of parametric t-SNE (van der Maaten, 2009), also used by Li et al. (2016) and Xu et al. (2020), utilizes a network consisting of a series of fully connected nodes with sigmoid activations between them. Xu et al. (2020) use three fully connected layers, with 500, 500, and 2,500 nodes, respectively, followed by a final fully connected layer with nodes (Fig. D.1). The latter architecture was used in the implementation of the present study with .

**Fig. D.1.**
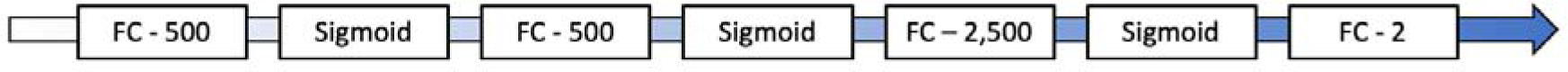
Illustration of the encoder of parametric t-SNE. FC: fully connected layer with the number of nodes indicated. Sigmoid: sigmoid activation.

In the original implementation, greedy layer-wise training is used to pretrain the encoders, followed by finetuning using t-SNE. In the present study, the encoders were trained directly using the principle of t-SNE. The same parameters were used as for the CNN encoders (batch size of 500, learning rate of 1e-4, and the same stopping criteria).

Two different types of feature representations (STFT, Appendix D.2.1; CWT, Appendix D.2.2) were used and the corresponding encoders had 1,823,502 and 2,726,502 parameters (all trainable), respectively.

### D.2. Features

In this work, two different time-frequency methods were used to generate features for parametric t-SNE: Short-time Fourier transforms (STFT) and continuous wavelet transforms (CWT). When producing transforms of the datasets described in Appendix A, the same randomizations were used so that all examples represented the same time interval across the different data representations and datasets.

#### D.2.1. STFT

STFT produce time series based on Fourier transforms for fixed sized windowed signals (Sejdić et al., 2009). All data were transformed using the STFT implementation in the SciPy library of Python (scipy.signal.stft), where each timestep represented 1 s. The default Hann window was used. The absolute of the amplitude values were taken. Frequencies up to 30 Hz (∼0.98–29.3Hz) were used and the total number of values was 630 per example (21 channels, 30 bands). The transform was first applied to whole continuous recordings before examples were extracted.

#### D.2.2. CWT

In their work using parametric t-SNE, Li et al. (2016) and Xu et al. (2020) use the Daubechies wavelet (Daubechies, 1988). This is a discrete wavelet where the resulting frequency bands, usually referred to as levels, have different time resolution. This has its benefits but can also make them harder to use and the duration of the examples will limit the frequency resolution. The frequency bands are relatively wide and sparse (Fig. D.2A); for some applications this may be a useful property by removal of unnecessary information and compressing the data. However, for other applications it may be detrimental, e.g., seizure activity may have a frequency evolution (Hirsch et al., 2021), and at least in theory, if the frequency bands are too wide and sparse then this may not be detected efficiently.

Since most signal examples of this study were of short duration, the continuous Ricker wavelet (also referred to as the Marr or Mexican hat wavelet) (Mallat, 2009) was also considered. The frequency bands can be made denser, thus providing a better frequency resolution (Fig. D.2B), but it comes at the expense of more overlapping of the bands. Tests were performed using the Python libraries pywt (Gregory et al., 2019) for the Daubechies wavelet and scipy.signal.cwt for the Ricker wavelet. For examples of 1 s duration, the Ricker wavelet showed a superior performance over the Daubechies wavelet when applying the original t-SNE on the resulting data (Fig. D.3A and D.3B).

Based on the above, the Ricker wavelet was thus chosen for this study, using widths 2, 3, 4, …, 30. The transform was applied to whole continuous recordings before extracting examples. For each channel and wavelet band of an example, the mean, the standard deviation, the minimum and the maximum value were calculated, in total producing 2436 values per example.

**Fig. D.2.**
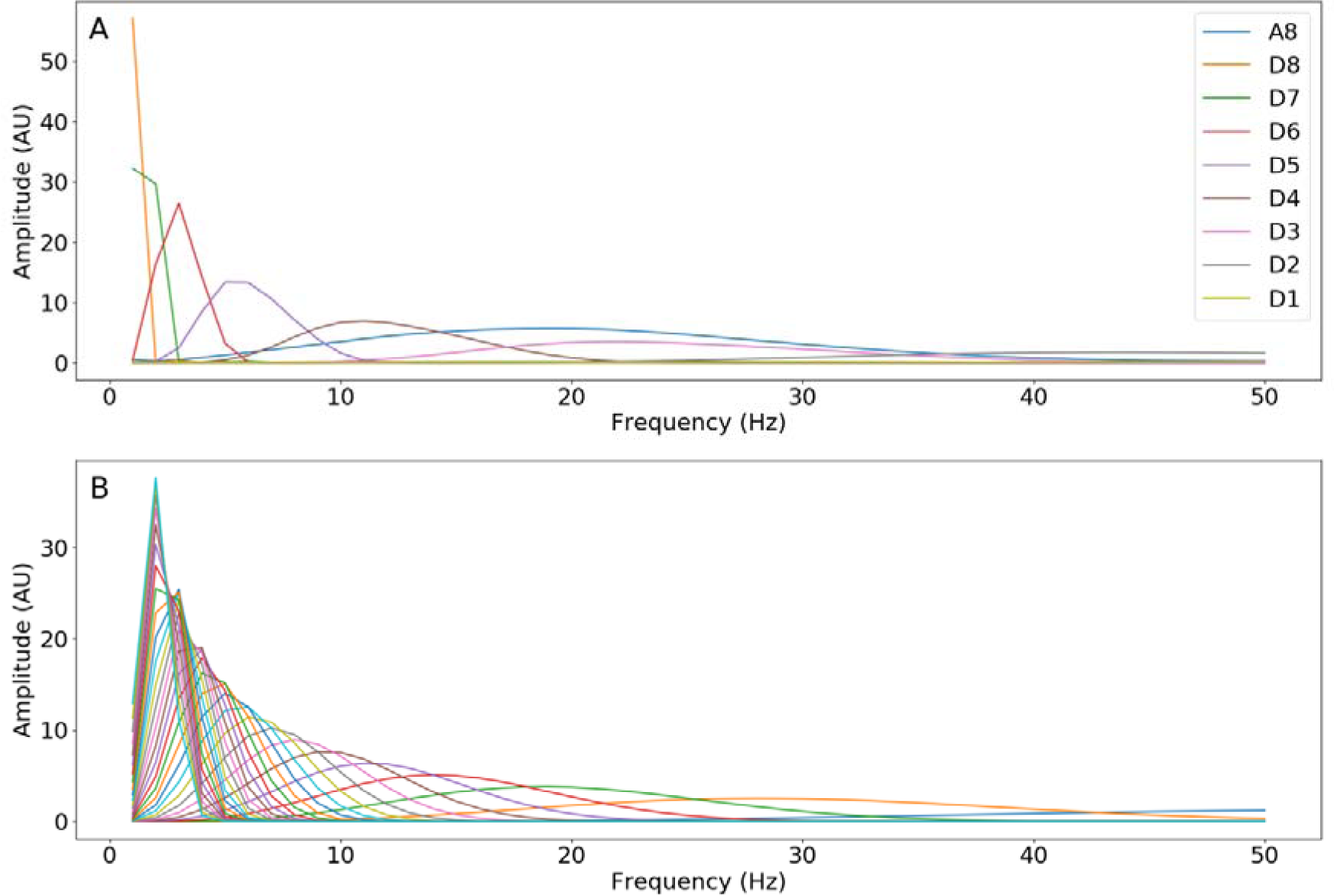
Frequency response for 1, 2, 3, …, 50 Hz. (A) The Daubechies wavelet (db4) with 8 levels. (B) The Ricker wavelet with widths 1, 2, 3, …, 30, where the maximum of each corresponding band has a frequency that decreases with increasing width.

**Fig. D.3.**
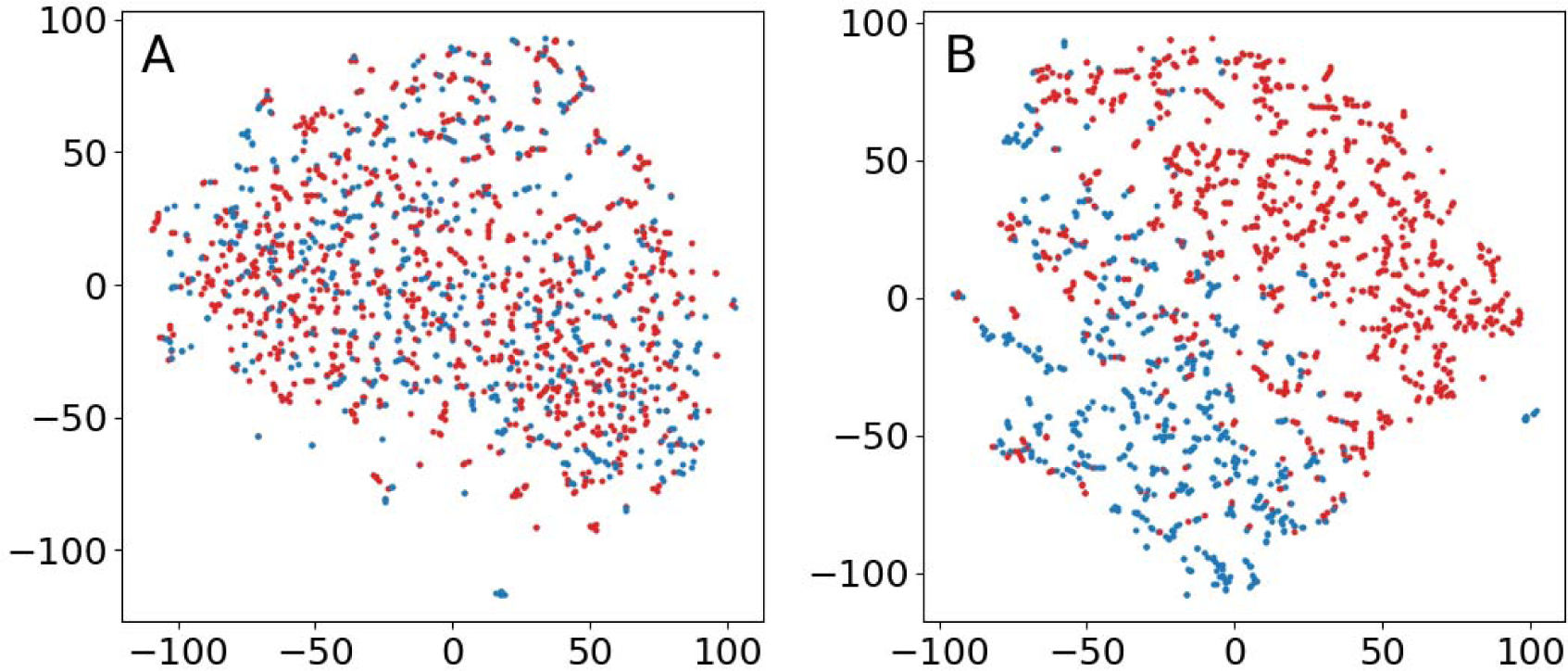
Results of t-SNE with perplexity 0.25 % for an EEG representing status epilepticus. Each dot represents 1 s and red dots indicate seizure activity. (A) The Daubechies wavelet (db4) using 6 levels. (B) The Ricker wavelet with widths 2, 3, 4, …, 30.

## Appendix E. Demonstration of sleep scoring using 30 s examples.

### E.1. Data

All but one of the extracted EEGs (1,000) were standard exams of about 20 minutes. The exception was a 12-hour recording from one subject containing sleep and wakefulness with a full range of sleep stages. This EEG has been used for some tests during the development of the method. The results presented here should therefore not be regarded as an evaluation test, but rather a demonstration of the method using examples of longer duration. It has therefore been placed in the appendix.

The EEG was divided into 30 s epochs and scored according to the AASM sleep scoring manual. An overview of the resulting distribution of sleep stages is given in Fig E.1A and E.1B. The 30 s duration was kept resulting in 1440 examples in total. The first 1080 examples were used for training and the last 360 examples was used for testing. Overviews of the distribution of sleep stages for training and test data are given in Fig E.1C and E.1D. (D) Distribution of sleep stages for test data.

**Fig. E.1.**
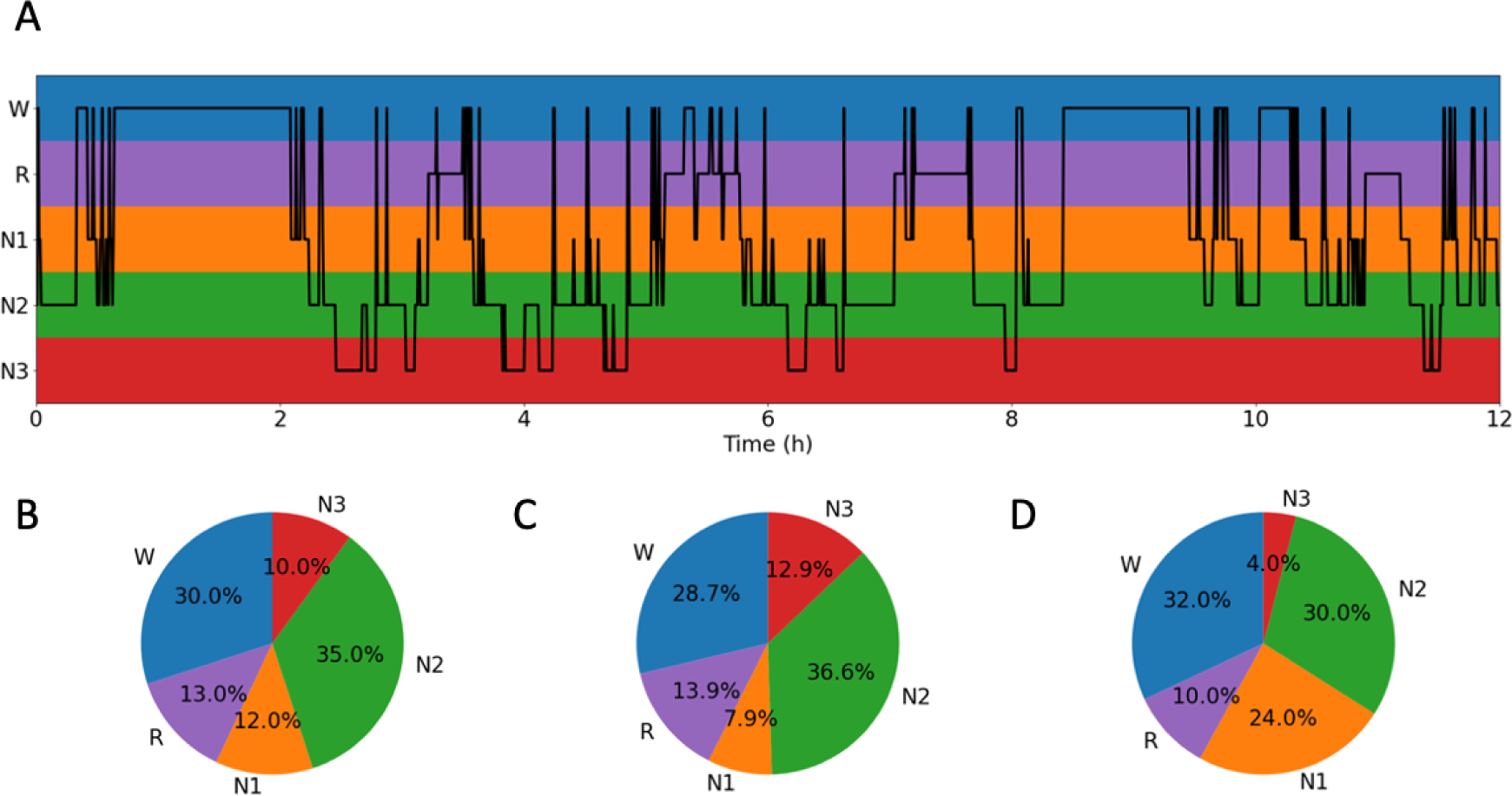
Sleep stages of 12-hour recording. (A) Overview of the sleep stages. (B) Distribution of the sleep stages for the whole recording. (C) Distribution of sleep stages for training data.

### E.2. Training encoders

Training was performed using a learning rate of 1e-4, a batch size of 500 and the number of neighbors and perplexity were set to 5. For the CNN encoders, due to memory constrains, to permit using the same batch size, the data were preprocessed: They were low pass filtered at 30 Hz and then downsampling to 62.5 Hz. It was also necessary to reduce the number of filters in the convolutional layers to half of what had been previously used. The time-frequency representations for the parametric t-SNE were generated as previously described (Appendix D). However, for the STFT features, for each channel and frequency band, the average values were calculated for the whole 30 s duration.

In addition, training was also performed for a CNN encoder using overlapping examples. A lower learning rate of 5e-6 was necessary for stable training. The training time for non-overlapping examples was approximately 20 hours and for overlapping examples and the lower learning rate, approximately 2 days and 14 hours.

### E.3. Results

Training with non-overlapping examples, resulted in similar kappa scores (Tab. E.1), the STFT encoder produced the highest scores, but there were no significant differences except for the training data and a linear kernel. In all instances, test scores were lower than training scores, indicating overfitting. A probable reason for this more apparent overfit, compared to the result in Section 3 of the article, is that the dataset used here was smaller and that there were five categories instead of two. Training clustering score for the CNN encoder was highest. Possibly due to outliers, the test clustering score for the STFT encoder was moderately high. The other clustering scores were low.

When training with overlapping examples, the results also showed overfitting. However, compared to the previous results for non-overlapping examples, it had decreased, and the test scores were significantly higher. For the linear kernel, the test score increased from 0.35 to 0.53, and for the RBF kernel it increased from 0.19 to 0.52. This comparison of course was not fair to the STFT and CWT encoders since no evaluations were performed using overlapping examples.

Visually, the results were in line with the quantitative measures (Fig. E.2). Separation and clustering were better for the STFT encoder and the CNN encoder when training with overlapping examples.

**Fig. E.2.**
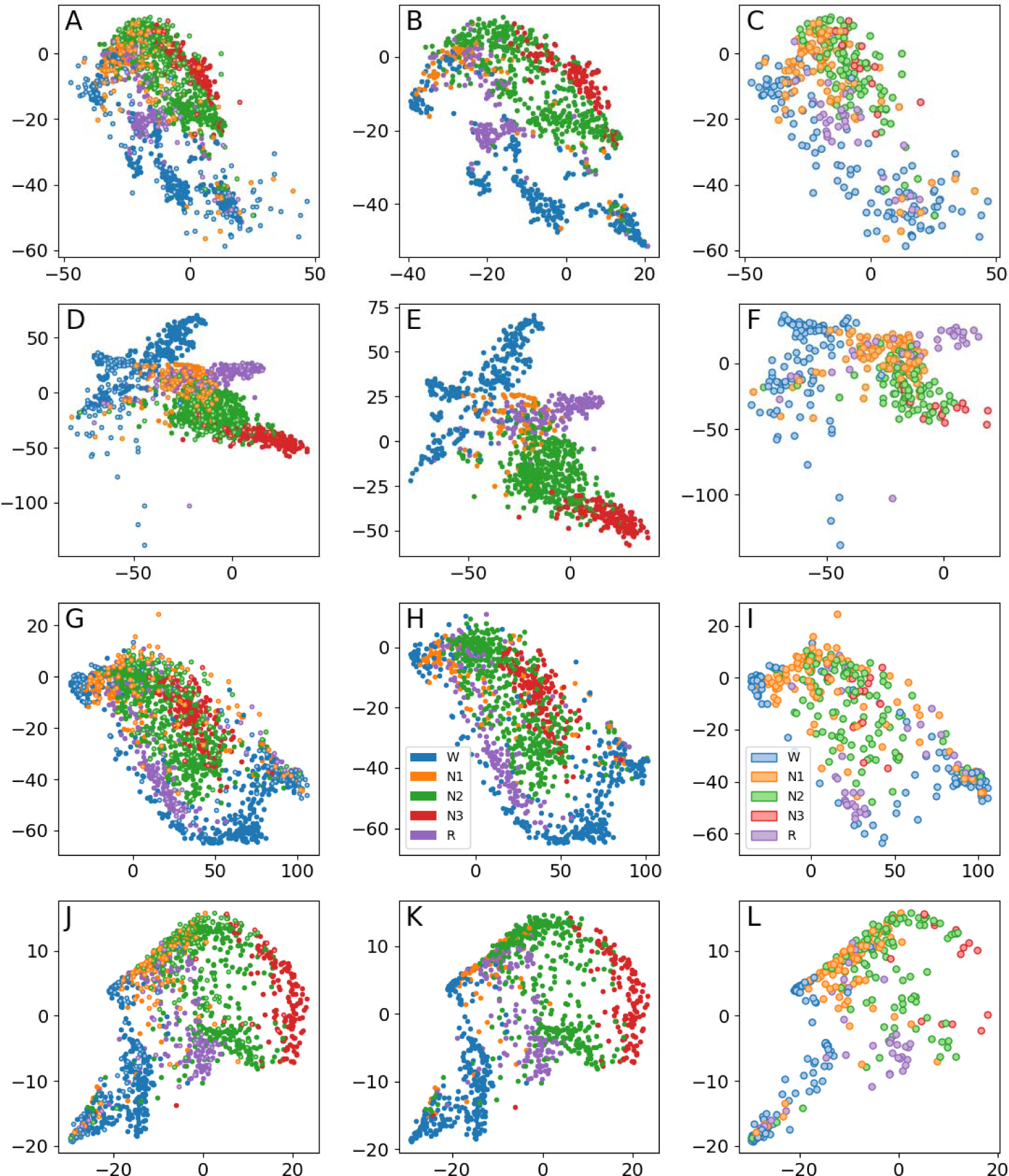
Low dimensional representations for the 12-hour dataset. Each dot represents 30 s of data. Blue = awake; orange = sleep stage N1; green: sleep stage N2; red: sleep stage N3; and purple: REM. First row (A–C): CNN encoder. Second row (D–F): STFT encoder. Third row (G–I): CWT encoder. Fourth row (J–L): CNN encoder using overlapping examples. First column (A, D, G, J): All data. Second column (B, E, H, K): Training data. Third column (C, F, I, L): Test data.

**Tab. E.1.** Quantitative measures for the 12-hour dataset. Significantly highest values in a column are in bold (chi-square test). *K* are kappa values for SVM, *L* indicate a linear kernel and *R* radialis basis function kernel; *C*. indicate the number of clusters by k-means.

| | $\kappa_L$-train | $\kappa_L$-test | $\kappa_R$-train | $\kappa_R$-test | $C$-train | $C$-test |
| --- | --- | --- | --- | --- | --- | --- |
| CNN | 0.55 | 0.35 | 0.85 | 0.19 | 12 | 2 |
| STFT | **0.77** | 0.51 | 0.96 | 0.33 | 2 | 6 |
| CWT | 0.36 | 0.30 | 0.92 | 0.30 | 2 | 2 |
| CNN* | 0.63 | **0.53** | 0.79 | **0.52** | 3 | 2 |

\*) Training with overlapping examples.

